# Human SLC46A2 is the dominant cGAMP importer in extracellular cGAMP-sensing macrophages and monocytes

**DOI:** 10.1101/2020.04.15.043299

**Authors:** Anthony F. Cordova, Christopher Ritchie, Volker Böhnert, Lingyin Li

## Abstract

Administration of exogenous CDNs to activate the cGAMP-STING pathway is a promising therapeutic strategy to unleash the full potential of cancer immunotherapy. This strategy mirrors the role of endogenous extracellular cGAMP, an immunotransmitter that is transferred from cancer cells to cGAMP-sensing cells in the host, promoting immunity. However, the CDN import mechanisms used by host cells within tumors remain unknown. Here we identified the protein SLC46A2 as the dominant cGAMP importer in primary human monocytes. Furthermore, we discovered that monocytes and M1-polarized macrophages directly sense tumor-derived extracellular cGAMP in murine tumors. Finally, we demonstrated that SLC46A2 is the dominant cGAMP importer in monocyte-derived macrophages. Together, we provide the first cellular and molecular mechanisms of cGAMP as an immunotransmitter, paving the way for effective STING pathway therapeutics.

## Introduction

Cancer immunotherapy has revolutionized the way in which cancer is treated, enabling physicians to now cure previously terminal diseases.^1^ Although most approved therapies target the adaptive immune system, activation of innate immune pathways is a prerequisite for these therapies to be effective. As such, there is growing interest in developing therapies that also target the innate immune system, such as the cytosolic double-stranded DNA (dsDNA) sensing cGAMP-STING pathway. Aberrant cytosolic dsDNA is a hallmark of cancer cells due to their intrinsic chromosomal instability, which is further enhanced by therapeutic ionizing radiation (IR).^2-3^ Detected as a danger signal, cytosolic dsDNA binds and activates the enzyme cyclic-GMP-AMP synthase (cGAS)^4^ to synthesize the cyclic dinucleotide (CDN) second messenger 2’3’-cyclic-GMP-AMP (cGAMP).^5–7^ cGAMP binds and activates the ER membrane protein Stimulator of Interferon Genes (STING). STING then activates TBK1, a kinase, and IRF3, a transcription factor, resulting in the expression of inflammatory cytokines.^8^ Of particular interest, the type I interferon (IFN-I) class of cytokines is necessary for cGAMP-mediated activation of T cells and effective anti-tumoral immunity.^9^

Although cancer cells constantly produce cGAMP,^10^ they often downregulate the canonical STING pathway and consequently do not produce high enough levels of IFN-I required for anti-cancer immunity.^11–13^ We recently discovered, however, that cancer cells secrete cGAMP into the tumor microenvironment as a soluble factor. Secreted cGAMP is an immunotransmitter that is internalized by cGAMP-sensing cells, leading to paracrine activation of the STING pathway and IFN-I production.^10^ This transfer of cGAMP is crucial for eliciting anti-tumoral immunity, as depletion of extracellular cGAMP in a murine breast tumor model abolished the curative effect of IR in a host-STING-dependent manner.^10^ Furthermore, intratumoral injections of cGAMP analogs demonstrated remarkable efficacy in murine models of cancer^14–15^ and are currently in phase I clinical trials for solid tumors (NCT02675439, NCT03172936, and NCT03937141).

As cGAMP has two negative charges, it is unable to passively diffuse across the cell membrane. Instead, extracellular cGAMP enters cells through cell-type-specific and species-specific transporters. We and others discovered that while human SLC19A1 is the dominant cGAMP transporter in some human cell lines, it is not the dominant transporter in all cell types.^16–17^ Additionally, there is no evidence that murine SLC19A1 is a cGAMP transporter. We recently discovered that the volume regulated chloride channel complex LRRC8A:C is the dominant cGAMP importer in primary human vasculature cells,^18^ while others reported that the LRRC8A:E complex is used by some murine cell types, including bone marrow-derived macrophages (BMDMs).^19^ However, the dominant cGAMP transporter used by primary human immune cells has not been identified. Additionally, it is unknown which specific immune cell types in the tumor microenvironment directly sense tumor-derived extracellular cGAMP and produce the IFN-Is necessary for anti-tumoral immunity. Identification of the cGAMP transporter used by these responder cells will be crucial in developing CDN-based immunotherapies, as STING activation in the improper cell type can lead to ineffective IFN-I production and impaired immunity^20^ or even immune cell death.^21–24^

Here, we identified the SLC46A family of solute carriers as novel cGAMP transporters and found that SLC46A2 is the dominant cGAMP transporter in human CD14^+^ monocytes. Additionally, we determined that intratumoral macrophages and NK cells directly sense endogenous extracellular cGAMP in murine tumors. Of particular interest, we found that M1-polarized, but not M2-polarized, intratumoral macrophages respond to tumor-derived extracellular cGAMP. Although murine M1-polarized macrophages do not express *Slc46a2*, we found that SLC46A2 is the dominant cGAMP importer in human monocyte-derived macrophages.

## Results

### CD14^+^ monocytes express high levels of the uncharacterized transporter SLC46A2

Monocytes and monocyte-derived cells, including macrophages and dendritic cells (DCs), play an important and complex role in the immune response to cancer. We and others previously found that primary human CD14^+^ monocytes are highly sensitive to extracellular cGAMP.^10, 25^ However, it is unknown how these cells import cGAMP. Although we previously characterized the reduced folic acid carrier SLC19A1 as an importer of cGAMP, we determined that SLC19A1 only plays a limited role as a cGAMP importer in CD14^+^ monocytes, as the SLC19A1 inhibitor methotrexate (MTX) had little effect on their extracellular cGAMP signaling in most healthy donors.^16^ Despite this, another inhibitor of SLC19A1, sulfasalazine (SSZ), strongly inhibited extracellular cGAMP signaling in CD14^+^ monocytes as determined by IRF3 phosphorylation (**Figure 1A**). We previously identified the LRRC8A channels as broadly expressed cGAMP transporters;^18^ however, SSZ is not known to inhibit LRRC8A channels. This suggests that SSZ is inhibiting an unknown cGAMP transporter in CD14^+^ monocytes. Furthermore, given that we previously reported that SSZ does not inhibit cGAMP signaling in the monocyte-derived U937 *SLC19A1^−/−^* cells,^16^ it appears that U937 cells do not express this unknown, SSZ-sensitive cGAMP transporter (**Figure 1B**). Thus, to identify additional cGAMP transporters, we compared expression levels of transmembrane transporters between CD14^+^ monocytes and U937 cells using published microarray data^26^ (**Figure 1C**). Of particular interest was *SLC46A2*, which encodes a transmembrane transporter that is highly expressed in CD14^+^ monocytes but not in U937 cells. Apart from one study that found that SLC46A2 is involved in the response to tracheal cytotoxin, relatively little is known about the function of SLC46A2.^27^ Given that SLC46A2 is closely related to the proton-coupled folic acid transporter SLC46A1, a known target of SSZ, we reasoned that SLC46A2 may be the cGAMP transporter in CD14^+^ monocytes.

**Figure 1.**
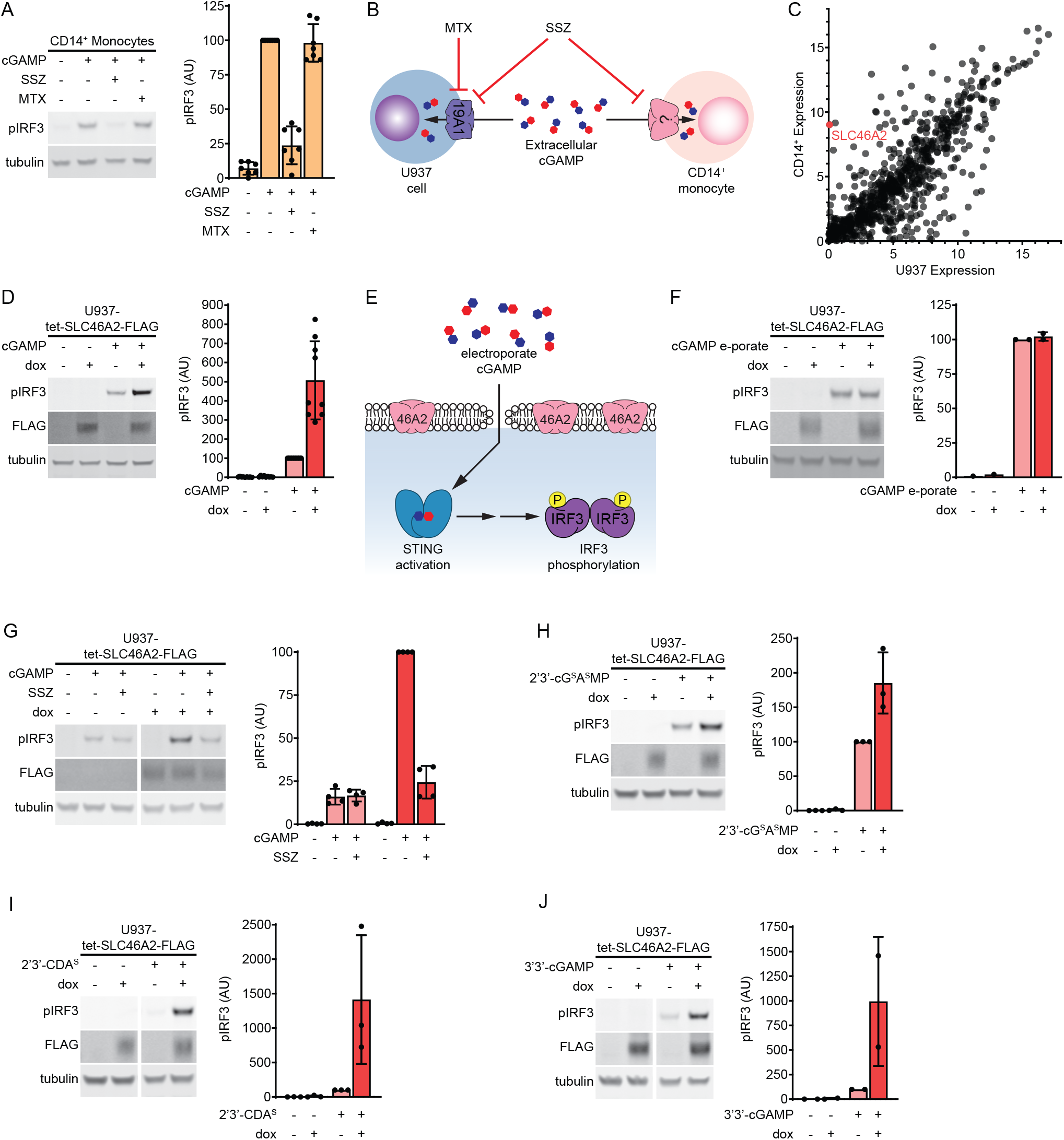
SLC46A2 is a cGAMP transporter. (**A**) Effect of sulfasalazine (SSZ) and methotrexate (MTX) on extracellular cGAMP signaling in CD14^+^ monocytes. Cells were pretreated with 1 mM SSZ or 500 μM MTX for 15 min and then treated with 50 μM cGAMP for 2 h. n = 7 individual donors. (**B**) Diagram illustrating effects of the SLC19A1 inhibitors SSZ and MTX on extracellular cGAMP signaling in U937 cells compared to CD14^+^ monocytes. (**C**) Microarray RNA expression levels of genes annotated as plasma membrane transmembrane transporters in U937 cells compared to CD14^+^ monocytes. (**D**) Effect of SLC46A2 overexpression on extracellular cGAMP signaling. U937-tet-SLC46A2-FLAG cells were induced with 1 μg/mL doxycycline (dox) for 24 h, then treated with 50 μM cGAMP for 2 h. n = 9 biological replicates. (**E**) Schematic illustrating how cGAMP electroporation bypasses cGAMP transporters. (**F**) Effect of SLC46A2 overexpression on intracellular cGAMP signaling. U937-tet-SLC46A2-FLAG cells were induced with 1 μg/mL dox for 24 h then electroporated with 100 nM cGAMP for 2 h. n = 2 biological replicates. (**G**) Effect of SSZ on SLC46A2 mediated cGAMP signaling. U937-tet-SLC46A2-FLAG cells were induced with 1 μg/mL dox for 24 h, then pretreated with 1 mM SSZ for 15 min before treatment with 50 μM cGAMP for 2 h. n = 4 biological replicates. (**H-J**) U937-tet-SLC46A2-FLAG cells were induced with 1 μg/mL dox for 24 h before treatment with either (**H**) 15 μM 2’3’-cG^S^A^S^MP, (**I**) 15 μM 2’3’-CDA^S^, or (**J**) 200 μM 3’3’-cGAMP for 2 h. n = 2-3 biological replicates. For (**A-J**), pIRF3 signal was normalized to tubulin signal, and data are shown as mean ± SD.

### SLC46A2 is a cGAMP transporter

In order to evaluate the potential role of human SLC46A2 protein as a cGAMP importer, we created a lentiviral vector encoding a C-terminally FLAG-tagged SLC46A2 under the control of a doxycycline inducible promoter (tet-SLC46A2-FLAG). This vector was transduced into U937 cells that had *SLC19A1* knocked out to reduce background cGAMP uptake (U937-tet-SLC46A2-FLAG). Using this cell line, we found that induction of SLC46A2-FLAG greatly increased the response to extracellular cGAMP (**Figure 1D**). While these data suggest that SLC46A2 is a cGAMP importer, it is possible that SLC46A2 is potentiating extracellular cGAMP signaling downstream of cGAMP import. To rule out this possibility, we evaluated the effect of SLC46A2 induction on the response to intracellular cGAMP that had been electroporated into cells (**Figure 1E**). In contrast to extracellular cGAMP signaling, SLC46A2 had no effect on intracellular cGAMP signaling, suggesting that SLC46A2 is a direct cGAMP importer (**Figure 1F**).

While multiple studies indicate that SLC46A2 localizes to the plasma membrane,^28–30^ a recent study found that SLC46A2 with a C-terminal EGFP tag localizes primarily to lysosomes.^27^ To verify that SLC46A2 is present on the plasma membrane where it can transport extracellular cGAMP, we inserted a FLAG tag into the first predicted extracellular loop of SLC46A2 (SLC46A2-exFLAG) (**Figure S1A**). We then transfected SLC46A2-exFLAG into HEK 293T cells and evaluated whether the FLAG tag was detectable on the cell surface through flow cytometry. As the cells were not permeabilized, only proteins on the cell surface should be detected. Indeed, we found that the SLC46A2-exFLAG cells were labeled with an anti-FLAG antibody but not with an antibody against the ubiquitous intracellular protein lamin A/C, indicating that SLC46A2 is present on the plasma membrane (**Figure S1B**).

We next evaluated the ability of SSZ to inhibit SLC46A2-mediated extracellular cGAMP signaling and found that SSZ treatment inhibited the effect of SLC46A2 induction (**Figure 1G**). Given that SLC19A1 and SLC46A2 are both inhibited by SSZ, we next tested whether other known inhibitors of SLC19A1 also inhibit SLC46A2. However, none of the competitive inhibitors of SLC19A1 (MTX, reduced folic acid (RFA), and oxidized folic acid (OFA)) significantly inhibited SLC46A2-mediated cGAMP signaling (**Figures S1C-E**). The SSZ metabolites 5-aminosalicylic acid (5-ASA) and sulfapyridine (SP) are thought to be the therapeutically active molecules in the treatment of inflammatory bowel disease and rheumatoid arthritis, respectively.^31^ Since the mechanisms of action of these metabolites are unknown, we tested whether 5-ASA or SP inhibited extracellular cGAMP signaling through SLC46A2. We found that 5-ASA did not reduce cGAMP signaling through SLC46A2 and SP only weakly reduced cGAMP signaling (**Figure S1F**), suggesting that these metabolites do not act by inhibiting SLC46A2.

### SLC46A2 selectively imports other CDNs

Given the chemical similarity across different CDNs, we tested whether SLC46A2 could import other CDNs in addition to cGAMP. Multiple synthetic CDNs, including 2’3’-bisphosphothioate-cGAMP (2’3’-cG^S^A^S^MP) and the investigative new drug 2’3’-bisphosphothioate-cyclic-di-AMP (2’3’-CDA^S^), have hydrolysis-resistant phosphothioate bonds in place of phophodiester backbones (**Figure S1G**). We found that induction of SLC46A2 increased the response to both 2’3’-cG^S^A^S^MP and 2’3’-CDA^S^, indicating that SLC46A2 can import substrates with a phosphothioate backbone (**Figures 1H-I**). While mammalian cGAMP contains both a 2’-5’ and a 3’-5’ phosphodiester bond, bacterial CDNs contain two 3’-5’ phosphodiester bonds (**Figure S1G**). Induction of SLC46A2 strongly increased the response to the bacterial CDN 3’3’-cGAMP (**Figure 1J**) and weakly increased the response to 3’3’-CDA (**Figure S1H**). Interestingly, SLC46A2 induction did not increase the response to another bacterial CDN, 3’3’-CDG (**Figure S1I**), demonstrating that SLC46A2 requires adenine rings to recognize CDNs but can tolerate diverse backbone linkages.

### SLC46A1 and SLC46A3 are CDN transporters

SLC46A2 is a member of the SLC46A solute transporter family, which also includes SLC46A1 and SLC46A3. SLC46A1 is the proton-coupled folic acid transporter, while the function of SLC46A3 remains largely uncharacterized. A previous study found that overexpression of SLC46A1 and SLC46A3 modestly enhanced STING pathway activation in response to extracellular cGAMP,^17^ suggesting that these proteins may also be cGAMP transporters in addition to SLC46A2. To confirm this, we generated U937 *SLC19A1^−/−^* cell lines expressing C-terminally FLAG-tagged SLC46A1 and SLC46A3 under the control of a doxycycline inducible promoter. Induction of SLC46A3 greatly increased the response to extracellular cGAMP (**Figure 2A**), while induction of SLC46A1 only modestly increased the response (**Figure 2B**). To confirm that the weak effect of SLC46A1 was not due to the FLAG tag interfering with transport, we also generated CRISPRa cell lines with guides targeting SLC46A1 and observed similar results (**Figure S2A**). The stronger effect of SLC46A3 induction on extracellular cGAMP signaling relative to SLC46A1 and SLC46A2 is likely due to higher expression levels of SLC46A3, as induction of the SLC46A3 transcript was approximately five-fold higher than induction of the SLC46A1 or SLC46A2 transcripts (**Figure S2B**). Using transcript levels to normalize the effect on extracellular cGAMP signaling across the three transporters, SLC46A2 increases the response by ~400%, SLC46A3 by ~100%, and SLC46A1 by ~50%. Like SLC46A2, both SLC46A1 and SLC46A3 were inhibited by SSZ, but not by MTX. However, the inhibitory effect of SSZ was significantly weaker against SLC46A3. Additionally, both RFA and OFA inhibited SLC46A1, while they only had a very minor inhibitory effect on SLC46A3. Electroporation of cGAMP into these cell lines abrogated the effect of doxycycline induction, demonstrating that these proteins are acting at the level of cGAMP import (**Figure S2C**). Taken together, these data indicate that SLC46A1, SLC46A2, and SLC46A3 are all cGAMP transporters, while SLC46A2 having the strongest activity.

**Figure 2.**
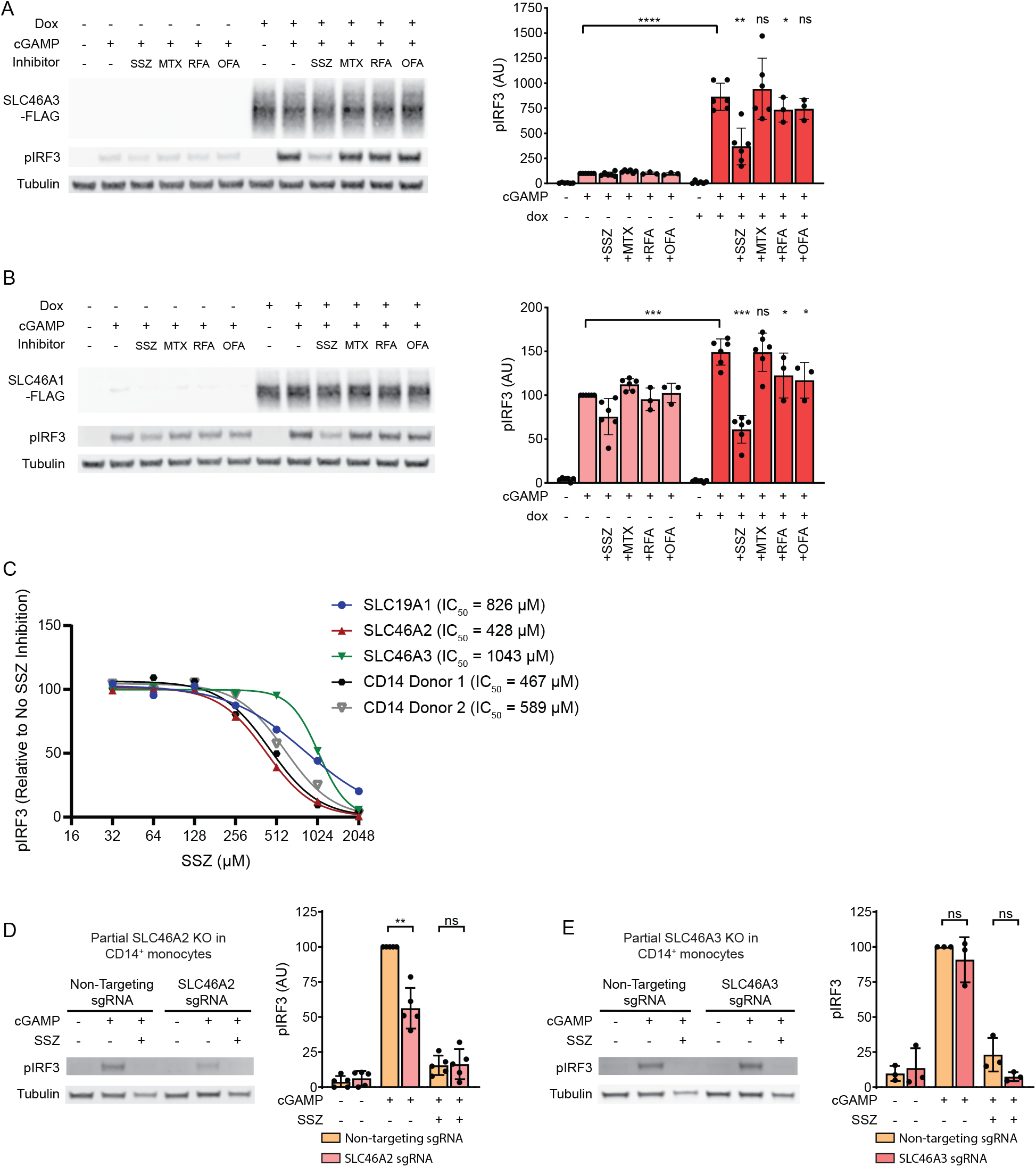
SLC46A2 is the dominant cGAMP importer in CD14^+^ monocytes. (**A-B**) U937-tet-SLC46A3-FLAG (**A**) or U937-tet-SLC46A1-FLAG (**B**) cells were induced with 1 μg/mL dox for 24 h. The cells were then pretreated with 1 mM SSZ, 500 μM MTX, 500 μM RFA, or 500 μM OFA for 15 min and then treated with 100 μM cGAMP for 90 min. n = 3 biological replicates. Data are shown as mean ± SD. (**C**) Dose-dependent inhibition of SSZ on SLC19A1, SLC46A2, and SLC46A3 compared to CD14^+^ monocytes. CD14^+^ monocytes and induced U937-tet-SLC-FLAG cells were pretreated with 32-2048 μM SSZ for 15 min before treatment with 50 μM cGAMP for 2 h. pIRF3 signal was normalized to tubulin, and then the data was normalized to fit an inhibition curve using a variable slope model, with the upper bound set at 100 and the lower bound set at 0. Donor 2 was normalized such that the highest concentration of SSZ corresponded to 100% inhibition, despite plateauing at 70% inhibition before normalization, suggesting the presence of an SSZ-insensitive minor transporter in this donor (see **Figure S2H**). n = 3 biological replicates for the cell lines, with only the mean shown for clarity. (**D**) Effect of partial CRISPR/Cas9 mediated knockout of SLC46A2 on extracellular cGAMP response in CD14^+^ monocytes. Freshly isolated CD14^+^ monocytes were electroporated with Cas9 RNPs targeting *SLC46A2*. 72 h after electroporation, cells were pretreated with 1 mM SSZ for 15 min and then treated with 50 μM cGAMP for 2 h. The percentage of *SLC46A2* gene knockout was estimated by TIDE analysis and ranged from 54%-80% (66% average). pIRF3 signal was normalized to tubulin signal, and data are shown as mean ± SD. n = 5 independent donors. (**E**) Effect of partial CRISPR/Cas9 mediated knockout of SLC46A3 on extracellular cGAMP response in CD14^+^ monocytes. Freshly isolated CD14^+^ monocytes were electroporated with Cas9 RNPs targeting *SLC46A3*. 72 h after electroporation, cells were pretreated with 1 mM SSZ for 15 min and then treated with 50 μM cGAMP for 2 h. The percentage of *SLC46A3* gene knockout was estimated by TIDE analysis and ranged from 48%-50% (49% average). pIRF3 signal was normalized to tubulin signal, and data are shown as mean ± SD. n = 3 independent donors.

We went on to determine if SLC46A1 and SLC46A3 can transport other CDNs. Consistent with SLC46A1 being a weak cGAMP transporter, overexpression of SLC46A1 only led to modest increases in response to 2’3’-cG^S^A^S^MP, 2’3’-CDA^S^ and 3’3’-cGAMP (**Figure S2D**). In contrast, overexpression of SLC46A3 led to large increases in the responses to 2’3’-cG^S^A^S^MP, 2’3’-CDA^S^, and 3’3’-cGAMP, similar to what we observed with SLC46A2 overexpression (**Figure S2E**). Given that both SLC46A2 and SLC46A3 appear to be strong CDN transporters, we evaluated the dose-response of these transporters over a range of CDN concentrations to determine their relative affinities for these CDNs. Across a range of cGAMP concentrations, SLC46A2 and SLC46A3 have nearly identical responses, suggesting they have similar affinities towards cGAMP (**Figure S2F**). When we looked across a range of 2’3’-CDA^S^ concentrations, however, we found that SLC46A2 was capable of responding to lower concentrations of 2’3’-CDA^S^ than SLC46A3, indicating that SLC46A2 is a higher affinity 2’3’-CDA^S^ transporter than SLC46A3 (**Figure S2G**).

### SLC46A2 is the dominant cGAMP importer in CD14^+^ monocytes

Given that SLC46A2 and SLC46A3 are both strong, SSZ-sensitive cGAMP transporters, we next sought to determine if either protein is the SSZ-sensitive cGAMP importer in primary CD14^+^ monocytes. We first compared inhibition of extracellular cGAMP signaling across a range of SSZ concentrations in doxycycline-induced U937-tet-SLC46A2-FLAG, U937-tet-SLC46A3-FLAG, and U937-tet-SLC19A1-FLAG cells. The inhibition curves were distinct for each protein, with SLC46A2 having the lowest IC_50_ (428 μM) and SLC46A3 having the highest IC_50_ (1043 μM). These SSZ inhibition curves were repeated on CD14^+^ monocytes from two independent donors, yielding inhibition curves very similar to the SLC46A2 curve, with IC_50_ values of 457 μM and 589 μM (**Figure 2C, S2H**). These data suggest that CD14^+^ monocytes use SLC46A2 to import cGAMP.

To validate these pharmacological results, we performed Cas9-mediated genetic knockout of SLC46A2 and SLC46A3 in primary CD14^+^ monocytes and then evaluated the response to extracellular cGAMP. We found that CD14^+^ monocytes with partial *SLC46A2* knockout (knockout efficiency 54-80%, average of 66%) had on average a 50% reduction in response to extracellular cGAMP (**Figure 2D**). In contrast, partial *SLC46A3* knockout (knockout efficiency 48-50%, average of 49%) had no significant effect on response to extracellular cGAMP (**Figure 2E**). Taken together, these data demonstrate that SLC46A2 is the dominant cGAMP importer in primary CD14^+^ monocytes.

### Intratumoral macrophages and monocytes directly sense tumor-derived extracellular cGAMP

Having demonstrated that primary human monocytes utilize SLC46A2 to import cGAMP, we next sought to characterize the role of monocytes and monocyte-derived cells as cGAMP-sensing cells within the tumor microenvironment. To this end, we established orthotopic 4T1-luciferase mammary tumors in BALB/c mice. As prior work has shown that ionizing radiation enhances the effect of extracellular cGAMP,^10^ the tumors were irradiated with 12 Gy once they reached 100 mm^3^. In order to specifically isolate the effects of extracellular cGAMP, we depleted extracellular cGAMP with intratumoral injections of soluble STING protein (neutralizing STING). A mutant version of STING that does not bind cGAMP (non-binding STING)^10^ was used as a control. Tumors were injected with 100 μM of STING protein 24 hours after irradiation, which is in excess of the predicted extracellular cGAMP concentration.^10^ After an additional 24 hours, tumors were extracted and analyzed by flow cytometry (**Figures 3A, S3**). For clarity, we will hereafter refer to tumors injected with non-binding STING as having extracellular cGAMP and those injected with neutralizing STING as not having extracellular cGAMP.

**Figure 3.**
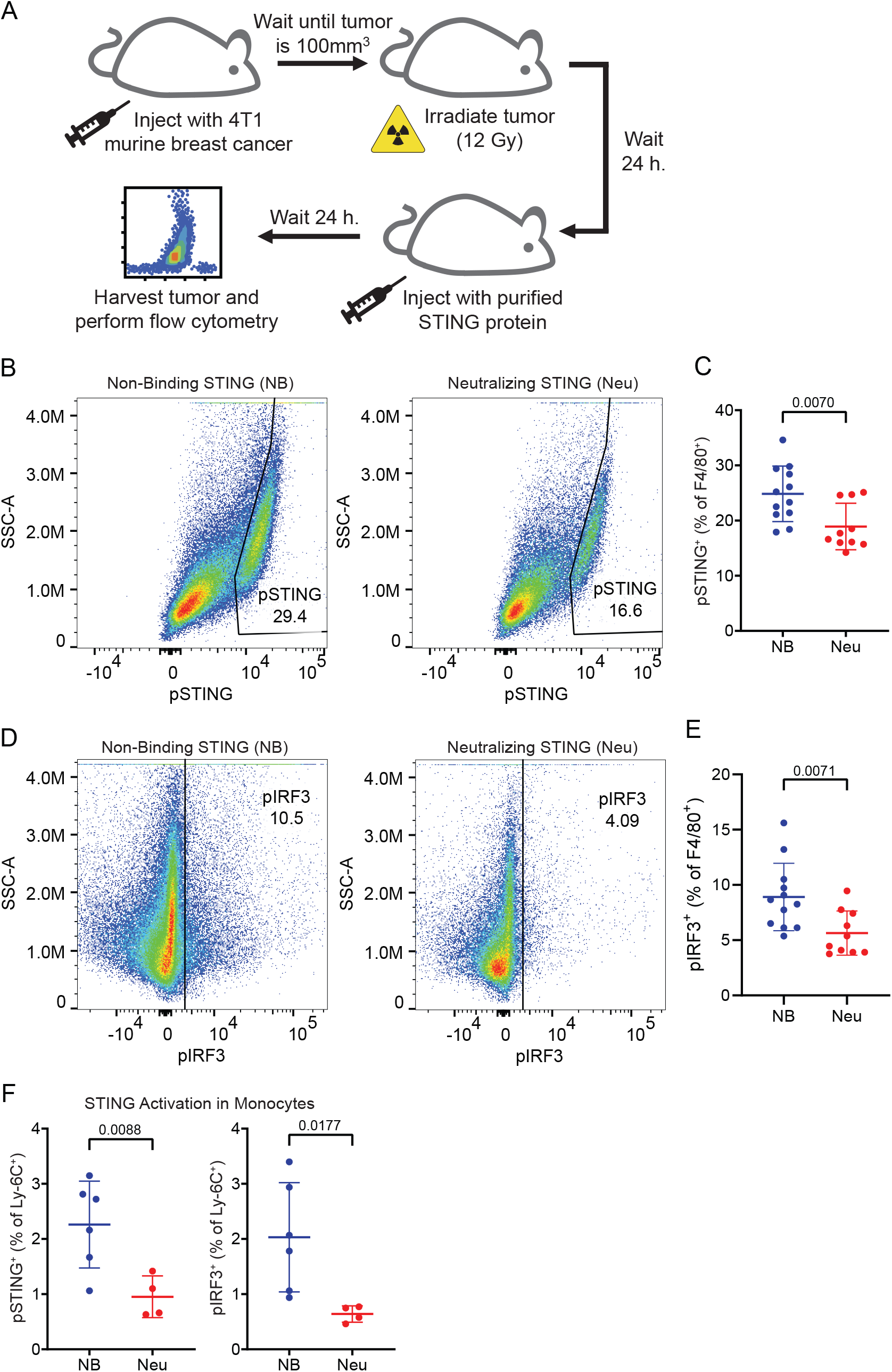
Intratumoral macrophages and monocytes directly sense tumor-derived extracellular cGAMP. (**A**) Experimental overview. BALB/c mice were injected with 50,000 4T1-Luciferase cells into the mammary fat pad. Once the tumors reached 100 mm^3^, the tumors were irradiated with 12 Gy. After 24 h, the tumors were injected with non-binding (NB) or neutralizing (Neu) STING. The mice were euthanized 24 h later and the tumors were extracted and prepared for flow cytometry. (**B-F**) Mice were included from 2 independent experiments as outlined in (**A**). Outliers were excluded using the ROUT method, and any tumors that were identified as outliers were removed from all analyses. n = 12 for NB STING (1 outlier removed) and n = 10 for Neu STING (2 outliers removed). Data is shown as the mean ± SD. p values were calculated by unpaired t test with Welch’s correction. (**B**) Representative flow cytometry plots identifying the pSTING^+^ populations as a percentage of F4/80^+^ macrophages in tumors from NB and Neu STING groups. (**C**) pSTING^+^ cells as a percentage of F4/80^+^ macrophages. Data is shown as the mean ± SD. p values were calculated by unpaired t test with Welch’s correction. (**D**) Representative flow cytometry plots identifying the pIRF3^+^ populations as a percentage of F4/80^+^ macrophages in tumors from NB and Neu STING groups. (**E**) pIRF3^+^ cells as a percentage of F4/80^+^ macrophages. Data is shown as the mean ± SD. p values were calculated by unpaired t test with Welch’s correction. (**F**) pSTING^+^ (left) and pIRF3^+^ (right) cells as a percentage of Ly-6C^+^ cells. Data are shown as the mean ± SD. p values were calculated by unpaired t test with Welch’s correction.

The presence of extracellular cGAMP had no effect on either total cell viability (**Figure S4A**) or overall immune cell infiltration (**Figure S4B**). Additionally, the presence of extracellular cGAMP did not significantly alter the immune composition of the tumors at this early timepoint, as there were no significant differences in the percentage of T cells, B cells, monocytes, macrophages, DCs, or NK cells within the tumor (**Figure S4C**). To identify the cell populations that directly respond to tumor-derived extracellular cGAMP, we probed the tumor samples for phosphorylated STING (pSTING) and phosphorylated IRF3 (pIRF3), which are both markers of STING pathway activation.^32–33^ Additionally, we probed for the IFN-Is interferon alpha (IFNα) and interferon beta (IFNβ), which are not specific for the STING pathway, but are functional consequences of its activation.

Because macrophages are a key cell type involved in the STING-mediated antitumoral immune response,^34–36^ we first analyzed intratumoral macrophages and their monocyte precursors. We found that macrophages had increased pSTING (**Figures 3B-C**) and pIRF3 (**Figures 3D-E**) signal in the presence of extracellular cGAMP, indicating that these cells internalize and respond to tumor-derived extracellular cGAMP. This was accompanied by increased IFN-I production (**Figure S4D**), demonstrating a functional immune response to extracellular cGAMP. Ly-6C^+^ cells, which are the murine monocytic precursors to macrophages,^37^ also had increased STING pathway activation in response to extracellular cGAMP (**Figure 3F**) but did not show an increase in IFN-I production (**Figure S4E**). Because STING activation typically occurs hours before IFN-I production, it is possible that Ly-6C^+^ cells internalized and responded to cGAMP more slowly than macrophages. Alternatively, it is possible that STING activation in Ly-6C^+^ cells resulted in their differentiation prior to IFN-I production.

### NK cells and T cells also directly sense tumor-derived extracellular cGAMP, but dendritic cells do not

In addition to macrophages and monocytes, we also profiled NK cells, which have been implicated in the cGAMP-mediated antitumoral immune response.^38–39^ We found that NK cells also had higher STING pathway activation (**Figure 4A**) and IFN-I production (**Figure 4B**) in the presence of tumor-derived extracellular cGAMP. NKG2D (also known as CD314) is expressed on mature NK cells and is upregulated after NK cell stimulation.^40–41^ NKG2D^Low^ NK cells showed an increase in STING pathway activation (**Figure 4C**) and IFN-I production (**Figure S5A**) in the presence of extracellular cGAMP, suggesting that the NKG2D^Low^ NK cells are direct cGAMP-sensing cells. NKG2D^High^ NK cells had an increase in IFN-I production (**Figure S5B**) without an increase in STING pathway activation (**Figure S5C**), suggesting that they are indirectly activated by extracellular cGAMP.

**Figure 4.**
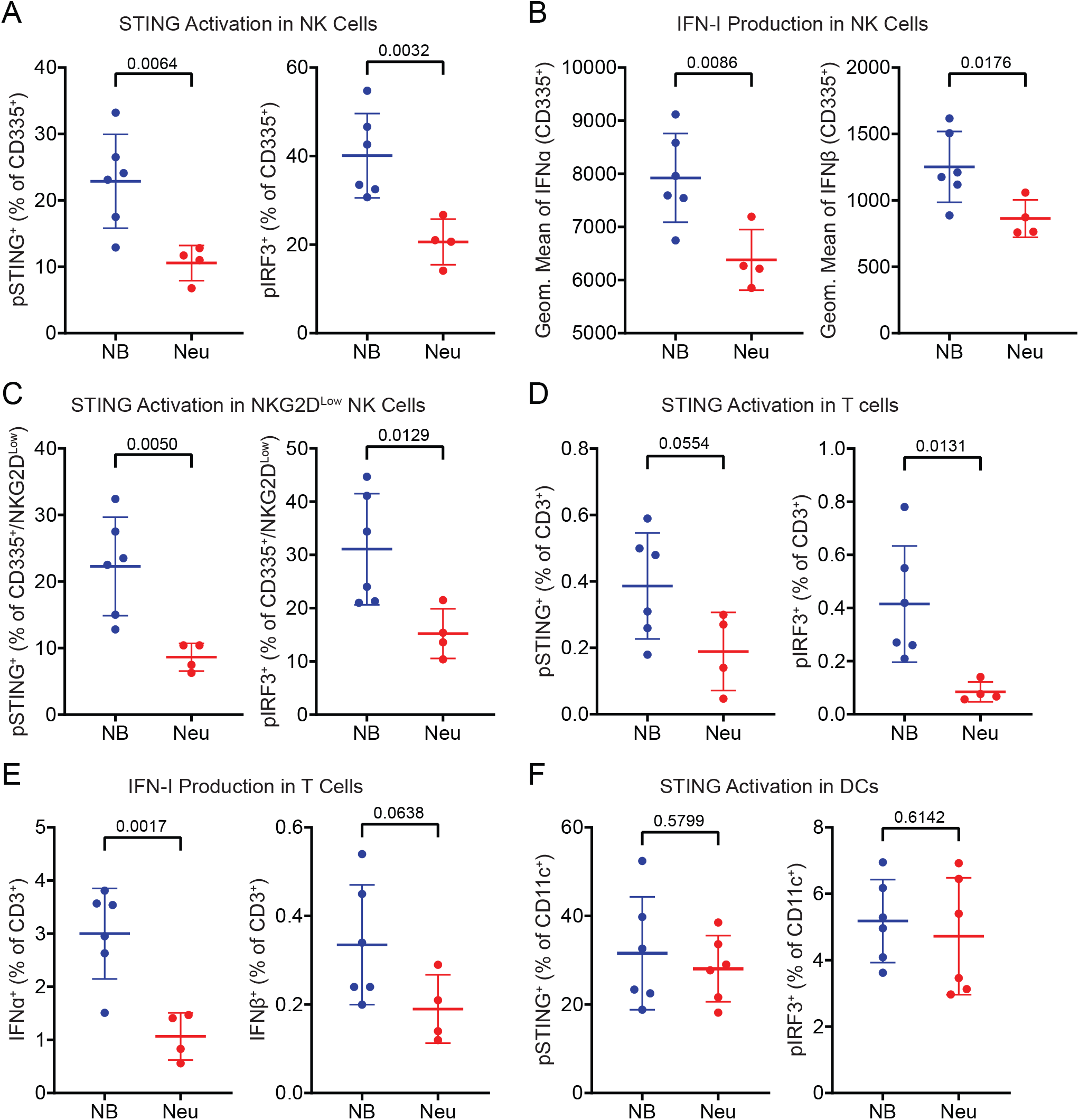
NK cells and T cells also directly sense tumor-derived extracellular cGAMP, but dendritic cells do not. (**A**) pSTING^+^ (left) and pIRF3^+^ (right) cells as a percentage of CD335^+^ NK cells. (**B**) Geometric mean of IFNα (left) and IFNβ (right) in CD335^+^ NK cells. (**C**) pSTING^+^ (left) and pIRF3^+^ (right) cells as a percentage of CD335^+^/NKG2D^Low^ NK cells. (**D**) pSTING^+^ (left) and pIRF3^+^ (right) cells as a percentage of CD3^+^ T cells. (**E**) IFNα^+^ (left) and IFNβ^+^ (right) cells as a percentage of CD3^+^ T cells. (**F**) pSTING^+^ cells (left) and pIRF3^+^ cells (right) as a percentage of CD11c^+^ DCs. (**A-F**) Data are shown as the mean ± SD. p values were calculated by unpaired t test with Welch’s correction.

Interestingly, a small percentage of T cells also had increased pSTING and pIRF3 (**Figure 4D**) signal in the presence of extracellular cGAMP, indicating that they are directly sensing extracellular cGAMP. These T cells also had higher IFN-I expression in the presence of extracellular cGAMP (**Figure 4E**), suggesting functional activation. These effects were primarily driven by CD4^+^ T cells (**Figures S5D-E**), although there may have been an increase in IFNβ production in CD8^+^ T cells (**Figures S5F-G**). However, only a small percentage of T cells sensed extracellular cGAMP, making it unlikely that T cells are a major cGAMP-sensing population.

Although there has been considerable evidence that DCs play a vital role in STING-mediated anti-tumoral immunity,^10, 42–43^ there was no difference in their STING pathway activation in the presence or absence of tumor-derived extracellular cGAMP (**Figure 4F**). This suggests that the role of DCs in STING-mediated anti-tumoral immunity is downstream of direct cGAMP-sensing cells. Likewise, there were no differences observed in B cells (**Figures S5H-I**). Together, these results demonstrate that only a specific subset of immune cells within the tumor directly sense tumor-derived extracellular cGAMP, with downstream effector cells being important for the subsequent immune response.

### M1-polarized macrophages are more sensitive to tumor-derived extracellular cGAMP

Tumor-associated macrophages comprise a wide range of cell states with varied and sometimes opposing roles,^44^ ranging from anti-tumoral M1 macrophages^45^ to pro-tumoral M2 macrophages.^46^ To distinguish between these macrophage states, we used the established cell surface marker CD206 (also known as MMR or MRC1), which is highly expressed in M2 macrophages (CD206^High^) and lowly expressed in M1 macrophages (CD206^Low^).^47–48^ We found that M1 (CD206^Low^) macrophages directly sensed tumor-derived extracellular cGAMP by activating their STING pathway (**Figures 5A-B**) and producing IFN-Is (**Figure 5C**), while M2 (CD206^High^) macrophages did not (**Figures 5D-E**). The number of M1 macrophages decreased as a percentage of total macrophages (**Figure S5J)**, possibly due to STING activation-induced death.^21^ It is also possible that the M1 population converted into M2 macrophages, as we observed a statistically insignificant increase in the absolute number of M2 macrophages in the tumor microenvironment (**Figures S5K-L**), despite there being prior evidence that cGAMP mediates the opposite conversion.^49^ Nevertheless, these data suggest that M1-polarized, but not M2-polarized, macrophages directly sense tumor-derived extracellular cGAMP and produce IFN-Is.

**Figure 5.**
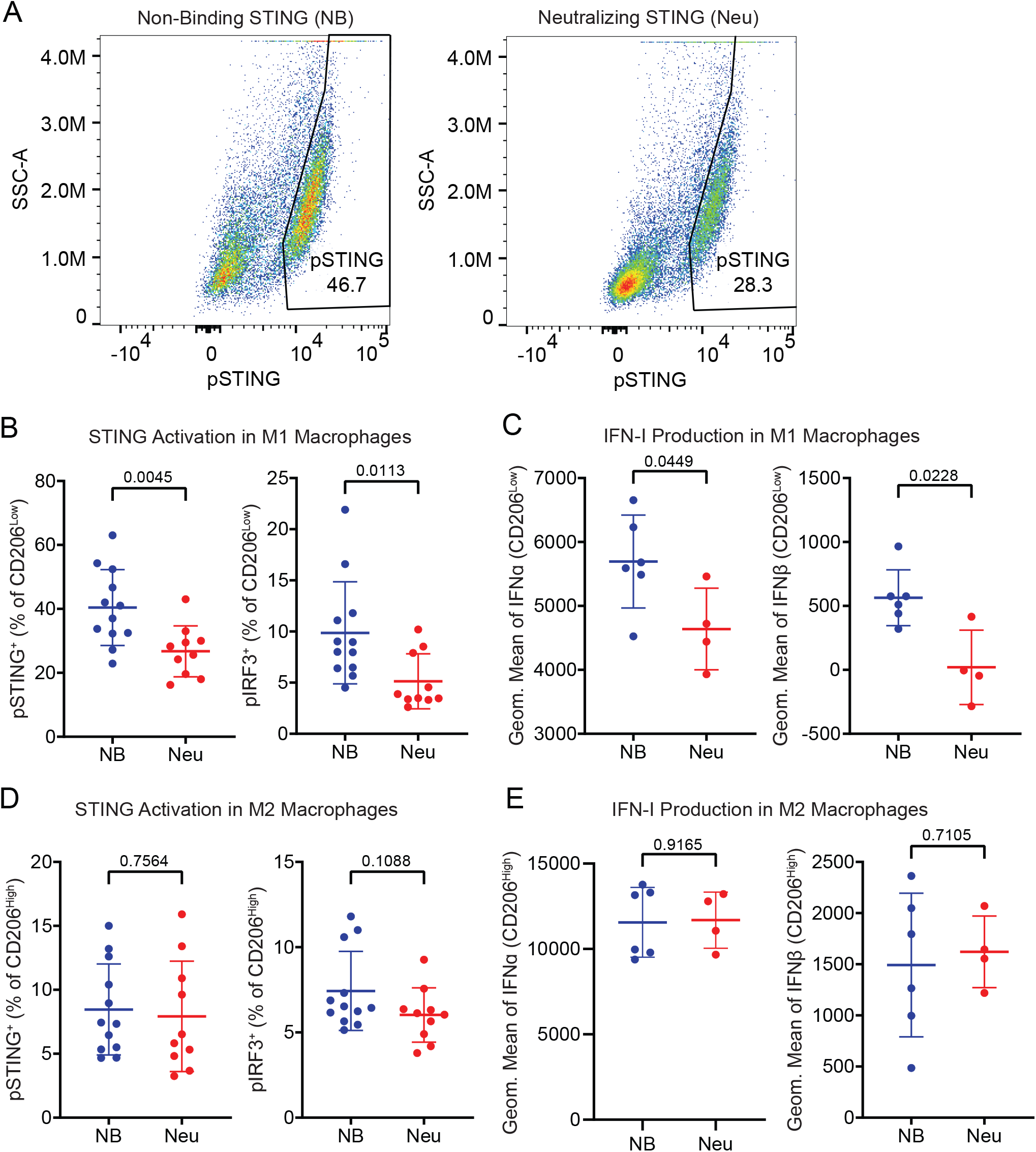
M1-polarized macrophages are more sensitive to tumor-derived extracellular cGAMP. (**A**) Representative flow cytometry plots identifying the pSTING^+^ populations as a percentage of F4/80^+^/CD206^Low^ M1 macrophages in tumors from NB and Neu STING groups. (**B**) pSTING^+^ (left) pIRF3^+^ (right) cells as a percentage of F4/80^+^/CD206^Low^ M1 macrophages. (**C**) Geometric mean of IFNα (left) and IFNβ (right) in F4/80^+^/CD206^Low^ M1 macrophages. (**D**) pSTING^+^ (left) and pIRF3^+^ (right) cells as a percentage of F4/80^+^/CD206^High^ M2 macrophages. (**E**) Geometric mean of IFNα (left) and IFNβ (right) in F4/80^+^/CD206^High^ M2 macrophages. (**A-F**) Data are shown as the mean ± SD. p values were calculated by unpaired t test with Welch’s correction.

### Intratumoral murine macrophages do not utilize SLC46A family members as cGAMP transporters

Having confirmed that monocyte-lineage cells, and in particular M1 macrophages, directly sense extracellular cGAMP in the tumor microenvironment, we next sought to identify the cGAMP transporters in these murine cells. Overexpression of murine mSLC19A1 did not affect the response to extracellular cGAMP in *SLC19A1^−/−^* U937 cells, indicating that mSLC19A1 is unlikely to be a cGAMP transporter (**Figure S6A**). In contrast, overexpression of murine mSLC46A2 strongly increased the response to extracellular cGAMP **(Figure 6A**) but did not increase the response to electroporated, intracellular cGAMP (**Figure S6B**), indicating that it is a cGAMP transporter. Overexpression of murine mSLC46A1 and mSLC46A3 also increased the response to extracellular cGAMP (**Figure 6B-C**, **S6C**), suggesting that these murine homologs are also cGAMP transporters. Although mSLC46A1 and mSLC46A3 were inhibited by SSZ, mSLC46A2 was not strongly inhibited by SSZ, in contrast to its human homolog. mSLC46A1 was inhibited by the folates RFA and OFA, while none of the transporters were inhibited by MTX. These data show that unlike SLC19A1, the ability of the SLC46A family members to import cGAMP is conserved between mice and humans. However, expression levels of the SLC46A transporters vary by cell types and species. In human immune cells, the *SLC46A2* transcript is highly expressed in monocytes and pre-DCs (**Figure S6D**). However, *Slc46a2* is poorly expressed in murine immune cells^26^ (**Figure S6E**). In line with this, we found that murine intratumoral macrophages do not express appreciable levels of *Slc46a2* transcript. However, murine intratumoral macrophages do express *Slc46a1* and *Slc46a3* (**Figure S6F)**.

**Figure 6.**
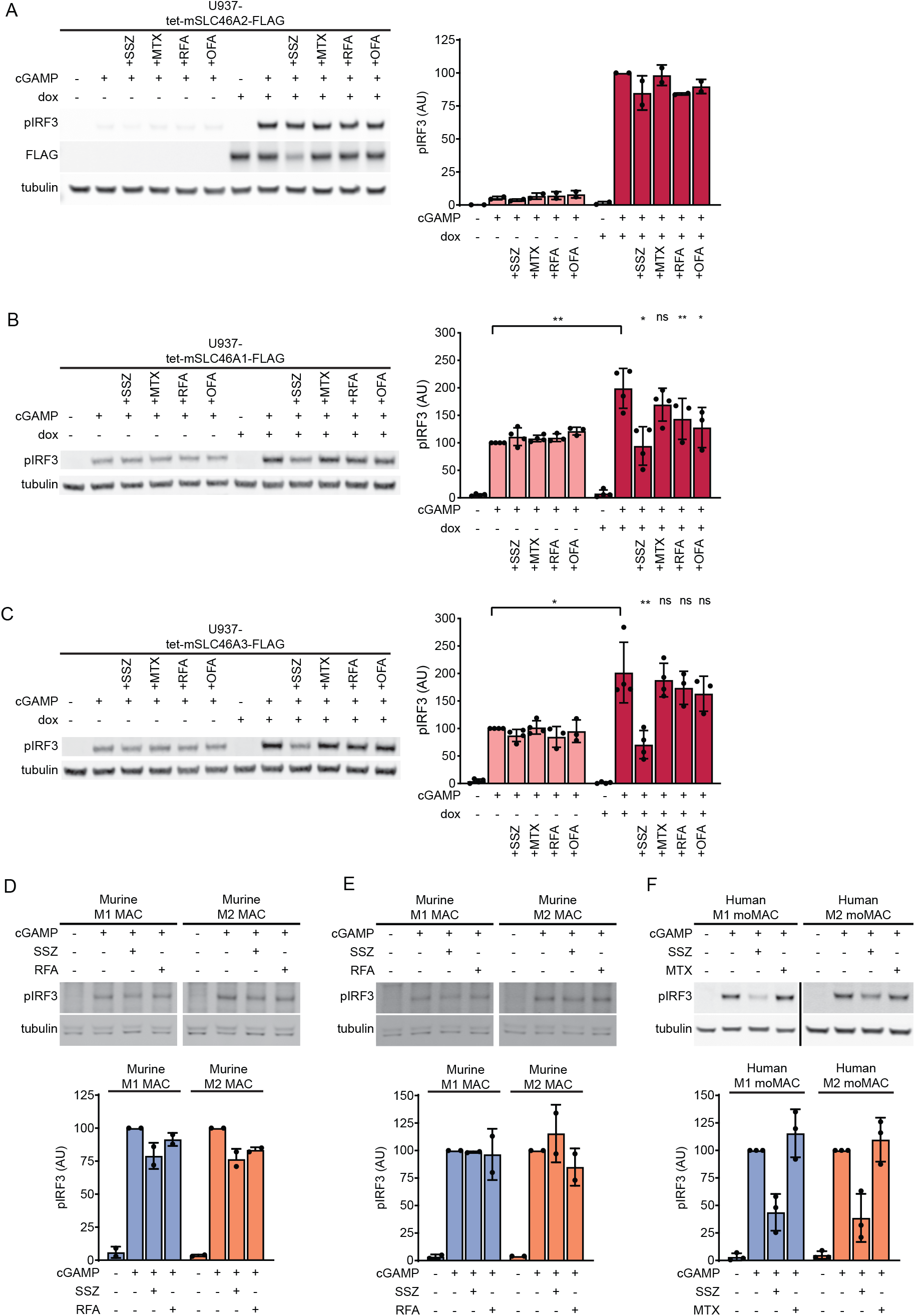
SLC46A2 is the dominant cGAMP importer in human, but not murine, macrophages. (**A**) Effect of mouse SLC46A2 on extracellular cGAMP signaling. U937-tet-mSLC46A2-FLAG cells were induced with 1 μg/mL dox for 24 h. The cells were then pretreated with 1 mM SSZ, 500 μM MTX, 500 μM RFA, or 500 μM OFA for 15 min and then treated with 50 μM cGAMP for 2 h. n = 2 biological replicates. (**B-C**) U937-tet-SLC46A1-FLAG (**B**) or U937-tet-SLC46A1-FLAG (**C**) cells were induced with 2 μg/mL dox for 48 h. The cells were then pretreated with 1 mM SSZ, 500 μM MTX, 500 μM RFA, or 500 μM OFA for 15 min and then treated with 100 μM cGAMP for 90 min. n = 3-4 biological replicates. Data are shown as mean ± SD. (**D-E**) BALB/c mice were injected with 50,000 4T1-Luciferase cells into the mammary fat pad. Once the tumors reached 100 mm^3^, half of the tumors were irradiated with 12 Gy. The mice were euthanized 48 h later and the tumors were extracted and prepared for FACS. 3-4 tumors were pooled into individual samples in order to increase the number of target cells. Cells were sorted into CD206^Low^ M1 and CD206^High^ M2 macrophages according to a gating scheme similar to Figure S2. The cells were pretreated with 1 mM SSZ or 500 μM RFA for 15 min and then treated with 50 μM cGAMP for 2 h. Non-irradiated samples are shown in (**D**) and irradiated samples are shown in (**E**). (**F**) Role of SLC46A2 in CD14^+^ monocyte-derived macrophages. Monocyte-derived M1 and M2 macrophages were treated with 50 μM cGAMP in the presence of either 1 mM SSZ or 500 μM MTX. n = 3 individual donors. For (**A-F**), pIRF3 signal was normalized to tubulin signal, and data are shown as mean ± SD.

To evaluate if mice utilize these transporters, we isolated intratumoral M1 and M2 macrophages from untreated and irradiated 4T1 tumors, as detailed above. Although both M1 and M2 intratumoral macrophages responded to extracellular cGAMP treatment, this response was only weakly inhibited by SSZ and RFA in the untreated tumors (**Figure 6D**) and not at all inhibited in the irradiated tumors (**Figure 6E**). Therefore, although mSLC46A1 and mSLC46A3 are capable of transporting cGAMP, they are not the dominant cGAMP transporters in murine intratumoral macrophages. Since the LRRC8A:E complex was recently identified as the primary cGAMP transporter in murine BMDMs,^19^ and intratumoral macrophages also express *Lrrc8a, Lrrc8c,* and a small amount of *Lrrc8e* (**Figure S6F**), it is possible that the LRRC8A channels are the primary cGAMP transporter in these cells as well, warranting future studies.

### SLC46A2 is the dominant cGAMP importer in human monocyte-derived macrophages

Given the species-specific usage of SLC46A2, we then endeavored to determine whether human M1 macrophages use SLC46A2 as a cGAMP transporter. Freshly isolated CD14^+^ monocytes were differentiated into either M1 macrophages or M2 macrophages using an established *in vitro* protocol^50^ and the effects of MTX and SSZ on extracellular cGAMP signaling were evaluated. MTX did not inhibit extracellular cGAMP signaling in either cell type, indicating that SLC19A1 is not a dominant cGAMP importer in human macrophages. In contrast, SSZ inhibited extracellular cGAMP signaling in both M1-and M2-polarized macrophages (**Figure 6F**). These data suggest that human monocyte-derived macrophages also utilize SLC46A2 as their dominant cGAMP importer.

## Discussion

In this study we demonstrated that murine M1 macrophages, in addition to NK cells, directly sense tumor-derived extracellular cGAMP, providing the first direct evidence of the extracellular cGAMP-STING-IRF3-IFN-I signaling cascade within tumors. Additionally, we identified SLC46A2 as the dominant cGAMP importer in human monocytes and monocyte-derived macrophages.

SLC46A2 is the third human cGAMP transporter identified after SLC19A1, a minor importer in CD14^+^ monocytes, and the LRRC8 channels, which are used by primary vascular cells. We hypothesize that different cell types in the tumor microenvironment express different levels of these and other cGAMP transporters. Since different transporters have distinct affinities toward extracellular cGAMP and varying transport kinetics, it is likely that both the local extracellular cGAMP concentration and transporter expression dictate which set of cells in the tumor microenvironment sense this immunotransmitter, and to what extent. For example, moderate concentrations of extracellular cGAMP might result in selective cGAMP import into IFN-I producing cells to promote immunity, whereas at higher concentrations cGAMP could also be imported by cells that die from cGAMP toxicity to prevent hyperinflammation.

The cell-type specific responses to extracellular cGAMP and other CDNs indicate that CDN-based therapeutics would be most effective when targeting the correct cell types to maximize the antitumoral immune response. Given that M1 macrophages produce high IFN-I levels in response to extracellular cGAMP signaling, optimizing therapeutics to specifically target their transporter SLC46A2 may result in more effective anticancer therapeutics. As the current CDN-based therapies in clinical trials are limited by STING-induced T cell toxicity,^25^ targeting cell-type specific importers could also reduce signaling in unwanted cell types. However, the species-specific usage of transporters tells a cautionary tale of testing CDN-based STING agonists in mice, despite mouse and human STING behaving similarly towards 2’3’-CDNs.

The origins of STING signaling are evolutionarily ancient, with STING homologs present in some bacteria.^51^ Throughout its evolution, STING signaling in different species has been fine-tuned to best suit the needs of that species. For example, while bacterial CDNs are strong activators of mouse STING, they only weakly activate human STING.^5, 52–53^ In addition, while mouse cGAS is sensitive to both short and long dsDNA, human cGAS is selectively activated by longer dsDNA.^54^ The observation that similar cell types in mice and humans use different cGAMP transporters is likely another example of species-specific divergence in STING signaling due to the different evolutionary pressures encountered by mice and humans.

Beyond the role of the STING pathway in anti-cancer immunity, it has previously been shown that colon-resident bacteria in a murine model of colitis promote STING activation and inflammation partially independent of cGAS, suggesting that host cells are able to import and respond to bacterial-synthesized CDNs.^55^ The bacterial CDNs 3’3’-cGAMP and 3’3’-CDA are associated with pathogenic bacteria,^56–58^ while 3’3’-CDG is produced by a wide variety of bacteria, including commensals.^59^ The ability of SLC46A2 and other CDN transporters^16, 18^ to selectively import certain CDNs (such as cGAMP and 3'3'-CDA) but not others (3'3'-CDG) suggests that CDN transporters could regulate how the immune system differentially responds to pathogenic and commensal bacteria.

In addition to being a hallmark of cancer, cytosolic dsDNA is also present in a wide variety of pathologies, including viral infections,^60–61^ myocardial infarction,^62^ autoimmune syndromes,^63–64^ pancreatitis,^65^ and aging.^66^ Since most cells export cGAMP as it is accumulates in the cytosol,^10^ it is likely that extracellular cGAMP plays a role in immune activation outside of cancer. Consequently, SLC46A2-bearing macrophages and their monocyte precursors could also be involved in sensing of extracellular cGAMP in these settings, and SLC46A2 inhibitors, such as FDA-approved SSZ, may alleviate excessive STING-mediated inflammation in these conditions.

## Acknowledgements

We thank all Li Lab members for their constructive comments and discussion through the course of this study. A.F.C. was supported by NIH 5T32GM736544 and 1F30CA250145. C.R. was supported by NIH 5T32GM007276. This work was supported by NIH DP2CA228044 (L.L). Cell sorting/flow cytometry analysis for this project was supported by the Stanford Shared FACS Facility.

## Author Contributions

A.F.C, C.R., V.B., and L.L. designed the study. A.F.C, C.R., and V.B. performed experiments. A.F.C, C.R., and L.L. wrote the manuscript. All authors discussed the findings and commented on the manuscript.

## Data and Materials Availability

All data are available in the main text or the supplementary materials.

## Methods

### Isolation of CD14^+^ Monocytes

Buffy coat (Stanford Blood Center) was diluted 1:3 with PBS supplemented with 2 mM EDTA. Diluted buffy coat was layered on top of 50% Percoll (GE Healthcare) containing 140 mM NaCl and centrifuged at 600 x g for 30 min. The separated PBMC layer was collected and washed once with PBS and once with RPMI. Following this, CD14^+^ cells were labeled using CD14 MicroBeads (Miltenyi Biotec) and isolated using a MACS LS Column on a MidiMACS Separator (Miltenyi Biotec) following the manufacturer’s instructions.

### Synthesis and purification of cGAMP

cGAMP was synthesized as previously described.^16^ To enzymatically synthesize cGAMP, 1 μM purified sscGAS was incubated with 50 mM Tris-HCl pH 7.4, 2 mM ATP, 2 mM GTP, 20 mM MgCl_2_, and 100 μg/mL herring testis DNA (Sigma) for 24 h. The reaction was then heated at 95°C for 3 min and filtered through a 3-kDa filter. cGAMP was purified from the reaction mixture using a PLRP-S polymeric reversed phase preparatory column (100 Å, 8 μm, 300 × 25 mm; Agilent Technologies) on a preparatory HPLC (1260 Infinity LC system; Agilent Technologies) connected to UV-vis detector (ProStar; Agilent Technologies) and fraction collector (440-LC; Agilent Technologies). The flow rate was set to 25 mL/min. The mobile phase consisted of 10 mM triethylammonium acetate in water and acetonitrile. The mobile phase started as 2% acetonitrile for first 5 min. Acetonitrile was then ramped up to 30% from 5-20 min, then to 90% from 20-22 min, maintained at 90% from 22-25 min, and then ramped down to 2% from 25-28 min. Fractions containing cGAMP were lyophilized and resuspended in water. The concentration was determined by measuring absorbance at 280 nm.

### Cell Culture

HEK 293T cells used for lentivirus generation and CD14^+^ cells used for CRISPR KO were maintained in DMEM with L-glutamine, 4.5g/L glucose and sodium pyruvate (Corning) supplemented with 10% FBS (Atlanta Biologicals) and 1% penicillin-streptomycin (GIBCO). U937 cells and all other CD14^+^ cells were maintained in RPMI (Corning) supplemented with 10% heat-inactivated FBS (Atlanta Biologicals) and 1% penicillin-streptomycin (GIBCO). 4T1-luciferase cells were a gift from Dr. Edward Graves^67^ and were maintained in RPMI (Cellgro) supplemented with 10% heat-inactivated FBS (Atlanta Biologicals) and 1% penicillin-streptomycin (GIBCO). All cells were maintained in a 5% CO_2_ incubator at 37 °C.

### Analysis of Microarray Data

Microarray data of RNA transcript expression levels in U937 cells and CD14^+^ monocytes from three donors was retrieved from Gene Expression Omnibus, accession GSE16076.^26^ For each microarray, background signal was subtracted from all probes so that the probe with the least signal was set to zero. Expression of transcripts in CD14^+^ monocytes was averaged across the three donors. Microarray probes targeting genes that were annotated in GeneOntology (accessed on 02-23-2020) as both transmembrane transporters (GO:0055085) and localized to plasma membrane (GO:0005886) were isolated to look for differential expression of transporters between U937 cells and CD14^+^ monocytes.

### Recombinant DNA

A plasmid containing the CDS of human SLC46A2 (pCMV-SPORT6-SLC46A2) was purchased from Harvard Plasmid Database. Plasmids containing the coding sequences of human SLC46A1 (pDONR221_SLC46A1) and SLC46A3 (pDONR221_SLC46A3) were purchased from Addgene. Custom plasmids (pTwist-CMV) containing the coding sequences of mouse *Slc19a1*, *Slc46a1*, *Slc46a2*, and *Slc46a3* were purchased from Twist Bioscience. To generate doxycycline inducible lentiviral plasmids, the transporter CDS was amplified from the appropriate plasmid using the primers listed in **Supplementary Table 1** and cloned into EcoRI/BamHI linearized pLVX-TetOne-FLAG-Hydro plasmid^18^ by isothermal Gibson assembly.^68^ To create a construct encoding SLC46A2 with an extracellular loop FLAG tag (pTwist-SLC46A2-exFLAG), a custom plasmid containing the codon optimized CDS of SLC46A2 (pTwist-SLC46A2) was linearized with AfeI and an oligonucleotide encoding a GGSG-linker flanked FLAG tag (**Supplementary Table 1**) was ligated into the cut site.

### Generation of Doxycycline Inducible Cell Lines

Lentiviral packaging plasmids (pHDM-G, pHDM-Hgmp2, pHDM-tat1b, and RC/CMV-rev1b) were purchased from Harvard Medical School. To generate lentivirus, 500 ng of lentiviral plasmid encoding doxycycline inducible transporters and 500 ng of each of the packaging plasmids were transfected into HEK 293T cells with FuGENE 6 transfection reagent (Promega). Cell supernatant was replaced 24 h after transfection and harvested after another 24 h. The lentivirus containing supernatant was passed through a 0.45 μm filter. To create the U937 *SLC19A1^−/−^* cell line,^16^ 1 mL filtered supernatant was supplemented with 8 mg/mL polybrene (Sigma Aldrich) and added to 1 × 10^5^ cells in a 24 well plate. Cells were spun at 1000 × g for 1 h, after which the virus containing media was removed and cells were resuspended in fresh media. After 48 h, cells were put under selection with the appropriate antibiotic alongside control cells (uninfected) until all control cells died.

### CDN Stimulation

U937 cells (0.5 × 10^6^ cells/mL) or freshly isolated CD14+ monocytes (1 × 10^6^ cells/mL) were treated with the indicated concentration of CDN for 2 h in a 5% CO_2_ incubator at 37 °C, unless otherwise indicated. Following treatments, cells were collected, lysed with Laemmli Sample Buffer, and run on SDS-PAGE gels for Western blot analysis.

### Electroporation of CDNs

U937 cells were pelleted and resuspended in nucleofector solution (90 mM Na_2_HPO_4_, 90 mM NaH_2_PO_4_, 5 mM KCl, 10 mM MgCl_2_, 10 mM sodium succinate) with the indicated CDN concentrations to a density of 1 × 10^6^ cells/mL. 100 uL cells were then transferred to a 0.2 cm electroporation cuvette and electroporated with program U-013 on a Nucleofector IIb device. Immediately after nucleofection, 500 uL media was added to cells. Cells were then transferred to a 24 well plate containing an additional 900 uL media and incubated in a 5% CO_2_ incubator at 37°C for 2 h. Following this, cells were collected, lysed with Laemmli Sample Buffer, and run on SDS-PAGE gels for Western blot analysis.

### Flow Cytometry to Determine SLC46A2 Localization

300 × 10^5^ HEK 293T cells were split onto 6 well plates and the next day were mock transfected or transfected with 1.5 ug pTwist-CMV-SLC46A2-exFLAG using 4.5 uL FuGene6 transfection reagent (Promega). 24 h after transfection, cells were trypsinized and transferred to 10 cm dishes 48 h after transfection, the cells were dissociated using PBS with 2 mM EDTA and then washed in PBS. The cells were stained with LIVE/DEAD Fixable Near-IR Dead Cell Stain (Invitrogen) for 30 min. The samples were then divided, with one half fixed and permeabilized with either eBioscience Foxp3/Transcription Factor Staining Buffer Set (Invitrogen), and the other half kept alive in PBS with 2% FBS. Samples were Fc-blocked for 10 min using TruStain FcX (BioLegend), and then stained for 45 min with mouse anti-Lamin A/C Alexa Fluor 488 Conjugate (Cell Signaling Technology) and rabbit anti-FLAG (DYKDDDDK Tag) (Cell Signaling Technology). The samples were then washed and stained with anti-rabbit Alexa Fluor 647 (Cell Signaling Technology) for 45 min. The samples were washed with PBS and then analyzed on a SH800S cell sorter (Sony Biotechnology).

### CRISPR KO of CD14^+^ Monocytes

Non-targeting, SLC46A2, and SLC46A3 sgRNAs were purchased from IDT and resuspended to 100 uM in TE buffer. Cas9 RNPs were formed by adding 8 uL of 61 uM Alt-R S.p. Cas9 Nuclease V3 (IDT) to 12 uL of 100 uM sgRNA and incubating for 10 min at room temperature. Freshly purified CD14^+^ monocytes were washed once with cold PBS, then resuspended in P3 Primary Cell nucleofector solution (Lonza) to a density of 10^7^ cells / 100 uL. 100 uL of resuspended monocytes were then added to the Cas9 RNPs, transferred to a nucleofection cuvette, and nucleofected using program CM-137 on a Nucleofector 4D device (Lonza). Electroporated cells were then transferred to a 6-well plate containing 2 mL of DMEM with 10% heat-inactivated FBS and 1% penicillin-streptomycin. 24 h after nucleofection, cells were pelleted and resuspended in 2 mL fresh media. 72 h after transfection, cells were used for CDN stimulation assays and genomic DNA was isolated to measure knockout efficiency. Knockout efficiency was determined by amplifying the region of genomic DNA surrounding sgRNA target sites (using the primers listed in **Supplementary Table 1**), performing Sanger sequencing, and using the sequencing trace to estimate knockout efficiency through TIDE analysis.^69^

### STING Expression and Purification

Wild-type (neutralizing) and R237A (non-binding) STING were expressed and purified using previously published methods.^10^ In brief, pTB146 His-SUMO-mSTING (residues 139-378) was expressed in Rosetta (DE3) pLysS competent cells (Sigma-Aldrich). Cells were grown in 2×YT medium with 100 μg/mL ampicillin until they reached an OD600 of 1. They were then induced with 0.75 mM IPTG at 16°C overnight. Cells were pelleted and resuspended in 50 mM Tris pH 7.5, 400 mM NaCl, 10 mM imidazole, 2 mM DTT, and protease inhibitors (cOmplete, EDTA-free protease inhibitor cocktail Roche). The cells were then flash frozen and thawed twice before sonication in order to lyse the cells. The lysate was then spun at 40,000 rpm at 4°C for 1 h. The supernatant was incubated with HisPur cobalt resin (Thermo Scientific) for 30 minutes at 4°C. The resin-bound protein was washed with 50 column volumes of 50 mM Tris pH 7.5, 150 mM NaCl, 2% triton X-114; 50 column volumes of 50 mM Tris pH 7.5, 1 M NaCl; and 20 column volumes of 50 mM Tris pH 7.5, 150 mM NaCl. Protein was eluted from resin with 600 mM imidazole in 50 mM Tris pH 7.5, 150 mM NaCl. Fractions containing His-SUMO-STING were pooled, concentrated, and dialyzed against 50 mM Tris pH 7.5, 150 mM NaCl while incubating with the SUMOlase His-ULP1 to remove the His-SUMO tag overnight. The solution was incubated with the HisPur cobalt resin again to remove the His-SUMO tag, and STING was collected from the flowthrough. Protein was dialyzed against 20 mM Tris pH 7.5, loaded onto a HitrapQ anion exchange column (GE Healthcare) using an Äkta FPLC (GE Healthcare), and eluted with a NaCl gradient. Fractions containing STING were pooled, buffer exchanged into PBS, and stored at − 80°C until use.

### Mouse Models

Mice were maintained at Stanford University in compliance with the Stanford University Institutional Animal Care and Use Committee (IACUC) regulations. All procedures were approved by the Stanford University Administrative Panel on Laboratory Animal Care (APLAC).

### Flow Cytometry Analysis of Tumors

7-9-week-old female BALB/c mice (Jackson Laboratories) were inoculated with 5 × 10^4^ 4T1-luciferase cells suspended in 50 μL of PBS. The cells were injected into the right fifth mammary fat pad. When tumor volume reached 100 ± 20 mm^3^, tumors were irradiated with 12 Gy using a 225 kVp cabinet X-ray irradiator with a 0.5 mm Cu filter (IC-250, Kimtron Inc.). Mice were anesthetized with a mixture of 80 mg/kg ketamine (VetaKet) and 5 mg/kg xylazine (AnaSed) prior to irradiation and were shielded with a 3.2 mm lead shield with 15 × 20 mm apertures to expose the tumors. Mice were then intratumorally injected with 100 μL of 100 μM neutralizing STING or non-binding STING 24 h after irradiation. Mice were euthanized 24 h later and the tumors were extracted. Following tumor extraction, the tumors were incubated in 10 mL of RPMI supplemented with 10% heat-inactivated FBS and 1% penicillin-streptomycin, as well as 20 μg/mL DNase I type IV (Millipore) and 1 mg/mL collagenase from Clostridium histolyticum (Sigma-Aldrich), at 37°C for 30 min. The samples were then passed through a 100 μm cell strainer (Sigma-Aldrich) to form a single-cell suspension. Red blood cells were lysed in 155 mM NH_4_Cl, 12 mM NaHCO_3_, and 0.1 mM EDTA for 5 min at room temperature. The samples designated for interferon detection were resuspended in 1 mL of RPMI supplemented with 10% heat-inactivated FBS and 1% penicillin-streptomycin and placed in a 5% CO_2_ incubator at 37°C for 1 h. 5 μg/mL Brefeldin A (BioLegend) was added to each sample, and they were incubated at 37°C for 5 additional hours before proceeding. All other samples proceeded directly to the live/dead stain after the red blood cell lysis. Samples were stained with LIVE/DEAD Fixable Blue Dead Cell Stain (Invitrogen) for 30 min. Samples were then fixed and permeabilized with either eBioscience Foxp3/Transcription Factor Staining Buffer Set (Invitrogen) or Fixation/Permeabilization Solution Kit (BD Biosciences). Samples were Fc-blocked for 10 min using TruStain fcX (BioLegend), and then stained for 1 h (see **Supplementary Table 2** for antibodies and dilutions). All samples were run on an Aurora analyzer (Cytek).

### FACS Sorting of Tumor Macrophages

7-9-week-old female BALB/c mice were injected with 50,000 4T1-Luciferase cells into the mammary fat pad. Once the tumors reached 100 mm^3^, the tumors were irradiated with 12 Gy. After 24 h, the tumors were injected with non-binding STING (this step was omitted for the experiment presented in **Figure 6D-E**). The mice were euthanized 24 h later and the tumors were extracted and prepared for FACS. 2-4 tumors were pooled into individual samples in order to increase the number of target cells. Following tumor extraction, the tumors were incubated in 10 mL of RPMI supplemented with 10% heat-inactivated FBS and 1% penicillin-streptomycin, as well as 20 μg/mL DNase I type IV (Millipore) and 1 mg/mL collagenase from Clostridium histolyticum (Sigma-Aldrich), at 37°C for 30 min. The samples were then passed through a 100 μm cell strainer (Sigma-Aldrich) to form a single-cell suspension. Red blood cells were lysed in 155 mM NH_4_Cl, 12 mM NaHCO_3_, and 0.1 mM EDTA for 5 min at room temperature. Samples were stained with LIVE/DEAD Fixable Blue Dead Cell Stain (Invitrogen) or LIVE/DEAD Fixable Near-IR Dead Cell Stain (Invitrogen) for 30 min, and then stained for 1 h (see **Supplementary Table 2** for antibodies and dilutions). Cells were sorted into CD206^High^ and CD206^Low^ macrophages using a FACSAria II (BD) cell sorter. The gating scheme for the sort was similar to the scheme presented in **Figure S3**.

### RNA-Seq

Sorted tumor cells were spun down and resuspended in 1 mL Trizol (Invitrogen) before being sent for RNA-seq. RNA-seq was performed by the Stanford Functional Genomics Facility. RNA was isolated using a guanidinium thiocyanate-phenol-chloroform extraction (TRIzol). Libraries were prepared using a Poly-A-enriched mRNA-Seq Library kit (KAPA) and were sequenced on a HiSeq 4000 (Illumina) using 2 × 75 bp paired-end reads. Demultiplexed reads were aligned to the GRCm38.p6 annotated mouse genome (GENCODE vM24) using STAR v2.7 in two-pass mode. Read counts for annotated genes were subsequently normalized to Transcripts Per Million (TPM).^70^

### Differentiation of CD14^+^ Monocytes

Freshly isolated CD14^+^ monocytes were differentiated into either M1 or M2 macrophages using a previously described phased protocol.^50^ In all three differentiation cases, CD14^+^ monocytes are seeded to a density of 3 × 10^5^ cells/mL in fresh RPMI media containing 10% heat-inactivated FBS and 1% penicillin-streptomycin on day 0; media was replaced on day 5; and a CDN stimulation experiment was performed on day 9. To differentiate into M1 macrophages, media was supplemented with 20 ng/mL GM-CSF on day 0, then supplemented with 20 ng/mL GM-CSF (PeproTech), 20 ng/mL IFN-γ (PeproTech), 20 ng/mL IL-6 (PeproTech), and 20 ng/mL LPS on day 5. To differentiate into M2 macrophages, media was supplemented with 20 ng/mL M-CSF (PeproTech) on day 0, then supplemented with 20 ng/mL M-CSF, 20 ng/mL IL-4, 20 ng/mL IL-6 (PeproTech), and 20 ng/mL IL-13 (PeproTech) on day 5.

### RT-qPCR

Total RNA was isolated from cells with TRIzol (Invitrogen) by following manufacturer’s protocol. To obtain cDNA, 20 uL RT reactions were setup containing 500 ng total RNA, 25 pmol oligo(dT)18, 25 pmol random hexamer primers, 0.5 mM dNTPs, 20 U RNaseOUT, 1x Maxima RT buffer, and 200 U Maxima RT (Thermo Scientific). RT reactions were incubated for 10 min at 25 °C, 15 min at 50 °C, then 5 min at 85 °C. To measure transcript levels, 10 uL qPCR reactions were setup containing 0.7 uL cDNA, 100 nM qPCR primers, and 1x AccuPower GreenStar master mix (Bioneer). To determine Ct values, reactions were run on a ViiA 7 Real-Time PCR System (Applied Biosystems) using the following program: ramp up to 50°C (1.6°C/s) and incubate for 2 min, ramp up to 95°C (1.6°C/s) and incubate for 10 min; then 40 cycles of ramp up to 95°C (1.6°C/s) and incubate for 15 s, ramp down to 60°C (1.6°C/s) and incubate for 1 min. Induced transcript levels were detected using primers that target the transcript’s 3’ UTR, and ACTB transcript levels were measured to normalize across samples (see **Supplementary Table 1**).

## Supplemental Figure Legends

**Figure S1.SLC46A2 is a cGAMP transporter.** (**A**) Predicted topology diagram of the first four transmembrane domains of SLC46A2 and insertion site of the extracellular FLAG tag. Note that only the first 165 amino acids (of 475) are depicted. Transmembrane helices were predicted with TMHMM v2.0. (**B**) HEK 293T cells were transfected with pTwist-CMV-SLC46A2-exFLAG. 72 h after transfection, cells were incubated with a live/dead stain and then half of the cells were fixed and permeabilized. The samples were then probed for lamin A/C and FLAG and analyzed by flow cytometry. Lamin A/C staining was included to ensure that the antibodies were not cell-permeable in the live condition. (**C-E**) Effects of the SLC19A1 inhibitors MTX, reduced folic acid (RFA), and oxidized folic acid (OFA) on SLC46A2-mediated cGAMP signaling. U937-tet-SLC46A2-FLAG cells were induced with 1 μg/mL dox for 24 h, then treated with 50 μM cGAMP in the presence of 500 μM of (**C**) MTX, (**D**) RFA, or (**E**) OFA for 2 h. n = 2-3 biological replicates. (**F**) Effects of the SSZ metabolites 5-aminosalicylic acid (5-ASA) and sulfapyridine (SP) on SLC46A2-mediated cGAMP signaling. U937-tet-SLC46A2-FLAG cells were induced with 1 μg/mL dox for 24 h, then pretreated with 1 mM 5-ASA or SP for 15 min before treatment with 50 μM cGAMP for 2 h. n = 2 biological replicates. (**G**) Chemical structures of synthetic and bacterial CDNs. (**H-I**) U937-tet-SLC46A2-FLAG cells were induced with 1 μg/mL dox for 24 h before treatment with either (**H**) 200 μM 3’3’-CDA or (**I**) 200 μM 3’3’-CDG for 2 h. n = 3 biological replicates. For (**C-I)**, pIRF3 signal was normalized to tubulin signal, and data are shown as mean ± SD.

**Figure S2. SLC46A2 is the dominant cGAMP importer in CD14^+^ monocytes.** (**A**) CRISPRa lines were generated in U937 *SLC19A1^-/-^* cells with either an off-target sgRNA or two different sgRNAs targeting SLC46A1. Cells were pretreated with 1 mm SSZ or 500 μM MTX for 15 and then treated with 100 μM cGAMP for 90 min. pIRF3 signal was normalized to tubulin signal, and data are shown as mean ± SD. (**B**) U937-tet-SLC46A1-FLAG or U937-tet-SLC46A3-FLAG cells were induced with 1 μg/mL dox for 24 h. Total RNA was purified, and induced transcript levels were measured through RT-qPCR. (**C**) Effect of SLC46A1 and SLC46A3 overexpression on intracellular cGAMP signaling. U937-tet-SLC46A1-FLAG and U937-tet-SLC46A3-FLAG cells were induced with 1 μg/mL dox for 24 h then electroporated with 100 nM cGAMP for 2 h. n = 2 biological replicates. (**D-E**) Effect of SLC46A1 and SLC46A3 overexpression on response to various CDNs. U937-tet-SLC46A1-FLAG (**D**) and U937-tet-SLC46A3-FLAG (**E**) cells were induced with 1 μg/mL dox for 24 h then treated with 25 μM 2’3’-cG^S^A^S^MP, 15 μM 2’3’-CDA^S^, or 200 μM 3’3’-cGAMP for 2 h. (**F**) Dose-response of SLC46A2 and SLC46A3 to cGAMP. U937-tet-SLC46A2-FLAG and U937-tet-SLC46A3-FLAG cells were induced with 1 μg/mL dox for 24 h then treated with 6.25-200 μM cGAMP for 2 h. (**G**) Dose-response of SLC46A2 and SLC46A3 to 2’3’-CDA^S^. U937-tet-SLC46A2-FLAG and U937-tet-SLC46A3-FLAG cells were induced with 1 μg/mL dox for 24 h then treated with 1.25-20 μM 2’3’-CDA^S^ for 2 h. (**H**) SSZ inhibition curves presented in **Figure 2C** prior to normalization of the upper and lower bounds, with donor 1 (left) and donor 2 (right) displayed separately for clarity.

**Figure S3. Gating scheme for flow cytometry analysis of tumors.** Immune cells were identified by CD45, and then further divided into immune subsets. T cells were identified by CD3, and then further divided into CD4^+^ and CD8^+^ T cells. B cells were identified by CD19, and DCs were identified by CD11c. Monocyte lineage cells were identified by Ly-6C. Macrophages were identified by F4/80, and then further divided into CD206^Low^ and CD206^High^ subsets. NK cells were identified by CD335, and then further divided into NKG2D^Low^ and NKG2D^High^. Gates were drawn by identifying clear populations or by comparing the experimental samples to unstained controls.

**Figure S4. Intratumoral macrophages and monocytes directly sense tumor-derived extracellular cGAMP.** (**A**) Live cells as a percentage of all singlets. (**B**) CD45^+^ immune cells as a percentage of all live cells. (**C**) Table displaying each cell population as a percentage of all live cells. No changes were statistically significant (p > 0.05). (**D**) Geometric mean of IFNα (left) and IFNβ (right) in F4/80^+^ macrophages. (**E**) IFNα^+^ (left) and IFNβ^+^ (right) cells as a percentage of Ly-6C^+^ cells. (**A**, **B**, **D**, **E**) Data are shown as the mean ± SD. p values were calculated by unpaired t test with Welch’s correction.

**Figure S5. M1-polarized macrophages, NK cells and T cells directly sense tumor-derived extracellular cGAMP, but dendritic cells do not.** (**A**) Geometric mean of IFNα (left) and IFNβ (right) in CD335^+^/NKG2D^Low^ NK cells. (**B**) Geometric mean of IFNα (left) and IFNβ (right) in CD335^+^/NKG2D^High^ NK cells. (**C**) pSTING^+^ (left) and pIRF3^+^ (right) cells as a percentage of CD335^+^/NKG2D^Low^ NK cells. (**D**) pSTING^+^ (left) and pIRF3^+^ (right) cells as a percentage of CD4^+^ T cells. (**E**) IFNα^+^ (left) and IFNβ^+^ (right) cells as a percentage of CD4^+^ T cells. (**F**) pSTING^+^ (left) and pIRF3^+^ (right) cells as a percentage of CD8^+^ T cells. (**G**) IFNα^+^ (left) and IFNβ^+^ (right) cells as a percentage of CD8^+^ T cells. (**H**) pSTING^+^ cells (left) and pIRF3^+^ cells (right) as a percentage of CD19^+^ B cells. (**I**) IFNα^+^ (left) and IFNβ^+^ (right) cells as a percentage of CD19^+^ B cells. (**J**) CD206^Low^ macrophages as a percentage of all live cells. (**K**) F4/80^+^/CD206^Low^ macrophages as a percentage of all live cells. (**L**) F4/80^+^/CD206^High^ macrophages as a percentage of all live cells. (**A-L**) Data are shown as the mean ± SD. p values were calculated by unpaired t test with Welch’s correction.

**Figure S6. SLC46A2 is the dominant cGAMP importer in human, but not murine, macrophages.** (**A**) Effect of mouse SLC19A1 on extracellular cGAMP signaling. U937-tet-mSLC19A1-FLAG cells were induced with 1 μg/mL dox for 24 h, then treated with 50 μM cGAMP. n = 3 biological replicates. Data are shown as mean ± SD. (**B**) Effect of mouse SLC46A2 on intracellular cGAMP signaling. U937-tet-mSLC46A2-FLAG cells were induced with 1 μg/mL dox for 24 h then electroporated with 100 nM cGAMP. n = 1 biological replicate. (**C**) U937-tet-mSLC46A1-FLAG or U937-tet-mSLC46A3-FLAG cells were induced with 2 μg/mL dox for 48 h. Total RNA was purified, and induced transcript levels were measured through RT-qPCR. (**D**-**E**) RNA-seq expression of SLC46A2 homologs in select (**D**) human and (**E**) mouse immune cells from Immgen. (**F**) BALB/c mice were injected with 50,000 4T1-Luciferase cells into the mammary fat pad. Once the tumors reached 100 mm^3^, the tumors were irradiated with 12 Gy. After 24 h, the tumors were injected with non-binding STING. The mice were euthanized 24 h later and the tumors were extracted and prepared for FACS. 2-3 tumors were pooled into individual samples in order to increase the number of target cells. Cells were sorted into CD206^High^ and CD206^Low^ macrophages according to a gating scheme similar to Figure S2 and then submitted for RNA-seq analysis. Expression values are displayed as transcripts per million (TPM). Data is presented as the mean of two samples per cell type, with the SD in parentheses.

## References

1. Sharma, P.; Allison, J. P., The future of immune checkpoint therapy. Science 2015, 348(6230), 56–61.

2. Harding, S. M.; Benci, J. L.; Irianto, J.; Discher, D. E.; Minn, A. J.; Greenberg, R. A., Mitotic progression following DNA damage enables pattern recognition within micronuclei. Nature 2017, 548 (7668), 466–470.

3. Mackenzie, K. J.; Carroll, P.; Martin, C. A.; Murina, O.; Fluteau, A.; Simpson, D. J.; Olova, N.; Sutcliffe, H.; Rainger, J. K.; Leitch, A.; Osborn, R. T.; Wheeler, A. P.; Nowotny, M.; Gilbert, N.; Chandra, T.; Reijns, M. A. M.; Jackson, A. P., cGAS surveillance of micronuclei links genome instability to innate immunity. Nature 2017, 548 (7668), 461–465.

4. Sun, L.; Wu, J.; Du, F.; Chen, X.; Chen, Z. J., Cyclic GMP-AMP synthase is a cytosolic DNA sensor that activates the type I interferon pathway. Science 2013, 339 (6121), 786–91.

5. Ablasser, A.; Goldeck, M.; Cavlar, T.; Deimling, T.; Witte, G.; Rohl, I.; Hopfner, K. P.; Ludwig, J.; Hornung, V., cGAS produces a 2'-5'-linked cyclic dinucleotide second messenger that activates STING. Nature 2013, 498 (7454), 380–4.

6. Gao, P.; Ascano, M.; Wu, Y.; Barchet, W.; Gaffney, B. L.; Zillinger, T.; Serganov, A. A.; Liu, Y.; Jones, R. A.; Hartmann, G.; Tuschl, T.; Patel, D. J., Cyclic [G(2',5')pA(3',5')p] is the metazoan second messenger produced by DNA-activated cyclic GMP-AMP synthase. Cell 2013, 153 (5), 1094–107.

7. Wu, J.; Sun, L.; Chen, X.; Du, F.; Shi, H.; Chen, C.; Chen, Z. J., Cyclic GMP-AMP is an endogenous second messenger in innate immune signaling by cytosolic DNA. Science 2013, 339 (6121), 826–30.

8. Ishikawa, H.; Ma, Z.; Barber, G. N., STING regulates intracellular DNA-mediated, type I interferon-dependent innate immunity. Nature 2009, 461 (7265), 788–92.

9. Deng, L.; Liang, H.; Xu, M.; Yang, X.; Burnette, B.; Arina, A.; Li, X. D.; Mauceri, H.; Beckett, M.; Darga, T.; Huang, X.; Gajewski, T. F.; Chen, Z. J.; Fu, Y. X.; Weichselbaum, R. R., STING-Dependent Cytosolic DNA Sensing Promotes Radiation-Induced Type I Interferon-Dependent Antitumor Immunity in Immunogenic Tumors. Immunity 2014, 41 (5), 843–52.

10. Carozza, J. A.; Böhnert, V.; Nguyen, K. C.; Skariah, G.; Shaw, K. E.; Brown, J. A.; Rafat, M.; von Eyben, R.; Graves, E. E.; Glenn, J. S.; Smith, M.; Li, L., Extracellular cGAMP is a cancer-cell-produced immunotransmitter involved in radiation-induced anticancer immunity. Nat Cancer 2020, 1, 184–196.

11. Bakhoum, S. F.; Ngo, B.; Laughney, A. M.; Cavallo, J. A.; Murphy, C. J.; Ly, P.; Shah, P.; Sriram, R. K.; Watkins, T. B. K.; Taunk, N. K.; Duran, M.; Pauli, C.; Shaw, C.; Chadalavada, K.; Rajasekhar, V. K.; Genovese, G.; Venkatesan, S.; Birkbak, N. J.; McGranahan, N.; Lundquist, M.; LaPlant, Q.; Healey, J. H.; Elemento, O.; Chung, C. H.; Lee, N. Y.; Imielenski, M.; Nanjangud, G.; Pe'er, D.; Cleveland, D. W.; Powell, S. N.; Lammerding, J.; Swanton, C.; Cantley, L. C., Chromosomal instability drives metastasis through a cytosolic DNA response. Nature 2018, 553 (7689), 467–472.

12. Xia, T.; Konno, H.; Ahn, J.; Barber, G. N., Deregulation of STING Signaling in Colorectal Carcinoma Constrains DNA Damage Responses and Correlates With Tumorigenesis. Cell Rep 2016, 14 (2), 282–97.

13. Ahn, J.; Xia, T.; Rabasa Capote, A.; Betancourt, D.; Barber, G. N., Extrinsic Phagocyte-Dependent STING Signaling Dictates the Immunogenicity of Dying Cells. Cancer Cell 2018, 33 (5), 862–873 e5.

14. Corrales, L.; Glickman, L. H.; McWhirter, S. M.; Kanne, D. B.; Sivick, K. E.; Katibah, G. E.; Woo, S. R.; Lemmens, E.; Banda, T.; Leong, J. J.; Metchette, K.; Dubensky, T. W.Jr.; Gajewski, T. F., Direct Activation of STING in the Tumor Microenvironment Leads to Potent and Systemic Tumor Regression and Immunity. Cell Rep 2015, 11 (7), 1018–30.

15. Curran, E.; Chen, X.; Corrales, L.; Kline, D. E.; Dubensky, T. W.Jr.,; Duttagupta, P.; Kortylewski, M.; Kline, J., STING Pathway Activation Stimulates Potent Immunity against Acute Myeloid Leukemia. Cell Rep 2016, 15 (11), 2357–66.

16. Ritchie, C.; Cordova, A. F.; Hess, G. T.; Bassik, M. C.; Li, L., SLC19A1 Is an Importer of the Immunotransmitter cGAMP. Mol Cell 2019, 75 (2), 372–381.

17. Luteijn, R. D.; Zaver, S. A.; Gowen, B. G.; Wyman, S. K.; Garelis, N. E.; Onia, L.; McWhirter, S. M.; Katibah, G. E.; Corn, J. E.; Woodward, J. J.; Raulet, D. H., SLC19A1 transports immunoreactive cyclic dinucleotides. Nature 2019, 573 (7774), 434–438.

18. Lahey, L. J.; Mardjuki, R. E.; Wen, X.; Hess, G. T.; Ritchie, C.; Carozza, J. A.; Bohnert, V.; Maduke, M.; Bassik, M. C.; Li, L., LRRC8A:C/E Heteromeric Channels Are Ubiquitous Transporters of cGAMP. Mol Cell 2020.

19. Zhou, C.; Chen, X.; Planells-Cases, R.; Chu, J.; Wang, L.; Cao, L.; Li, Z.; Lopez-Cayuqueo, K. I.; Xie, Y.; Ye, S.; Wang, X.; Ullrich, F.; Ma, S.; Fang, Y.; Zhang, X.; Qian, Z.; Liang, X.; Cai, S. Q.; Jiang, Z.; Zhou, D.; Leng, Q.; Xiao, T. S.; Lan, K.; Yang, J.; Li, H.; Peng, C.; Qiu, Z.; Jentsch, T. J.; Xiao, H., Transfer of cGAMP into Bystander Cells via LRRC8 Volume-Regulated Anion Channels Augments STING-Mediated Interferon Responses and Anti-viral Immunity. Immunity 2020, 52 (5), 767–781 e6.

20. Wang, J.; Li, P.; Yu, Y.; Fu, Y.; Jiang, H.; Lu, M.; Sun, Z.; Jiang, S.; Lu, L.; Wu, M. X., Pulmonary surfactant-biomimetic nanoparticles potentiate heterosubtypic influenza immunity. Science 2020, 367 (6480).

21. Gaidt, M. M.; Ebert, T. S.; Chauhan, D.; Ramshorn, K.; Pinci, F.; Zuber, S.; O'Duill, F.; Schmid-Burgk, J. L.; Hoss, F.; Buhmann, R.; Wittmann, G.; Latz, E.; Subklewe, M.; Hornung, V., The DNA Inflammasome in Human Myeloid Cells Is Initiated by a STING-Cell Death Program Upstream of NLRP3. Cell 2017, 171 (5), 1110–1124 e18.

22. Gulen, M. F.; Koch, U.; Haag, S. M.; Schuler, F.; Apetoh, L.; Villunger, A.; Radtke, F.; Ablasser, A., Signalling strength determines proapoptotic functions of STING. Nat Commun 2017, 8 (1), 427.

23. Cerboni, S.; Jeremiah, N.; Gentili, M.; Gehrmann, U.; Conrad, C.; Stolzenberg, M. C.; Picard, C.; Neven, B.; Fischer, A.; Amigorena, S.; Rieux-Laucat, F.; Manel, N., Intrinsic antiproliferative activity of the innate sensor STING in T lymphocytes. J Exp Med 2017, 214 (6), 1769–1785.

24. Larkin, B.; Ilyukha, V.; Sorokin, M.; Buzdin, A.; Vannier, E.; Poltorak, A., Cutting Edge: Activation of STING in T Cells Induces Type I IFN Responses and Cell Death. J Immunol 2017, 199 (2), 397–402.

25. Sivick, K. E.; Desbien, A. L.; Glickman, L. H.; Reiner, G. L.; Corrales, L.; Surh, N. H.; Hudson, T. E.; Vu, U. T.; Francica, B. J.; Banda, T.; Katibah, G. E.; Kanne, D. B.; Leong, J. J.; Metchette, K.; Bruml, J. R.; Ndubaku, C. O.; McKenna, J. M.; Feng, Y.; Zheng, L.; Bender, S. L.; Cho, C. Y.; Leong, M. L.; van Elsas, A.; Dubensky, T. W., Jr.; McWhirter, S. M., Magnitude of Therapeutic STING Activation Determines CD8(+) T Cell-Mediated Anti-tumor Immunity. Cell Rep 2018, 25 (11), 3074–3085 e5.

26. Gebhard, C.; Benner, C.; Ehrich, M.; Schwarzfischer, L.; Schilling, E.; Klug, M.; Dietmaier, W.; Thiede, C.; Holler, E.; Andreesen, R.; Rehli, M., General transcription factor binding at CpG islands in normal cells correlates with resistance to de novo DNA methylation in cancer cells. Cancer Res 2010, 70 (4), 1398–407.

27. Paik, D.; Monahan, A.; Caffrey, D. R.; Elling, R.; Goldman, W. E.; Silverman, N., SLC46 Family Transporters Facilitate Cytosolic Innate Immune Recognition of Monomeric Peptidoglycans. J Immunol 2017, 199 (1), 263–270.

28. Kim, M. G.; Flomerfelt, F. A.; Lee, K. N.; Chen, C.; Schwartz, R. H., A putative 12 transmembrane domain cotransporter expressed in thymic cortical epithelial cells. J Immunol 2000, 164 (6), 3185–92.

29. Ahn, S.; Lee, G.; Yang, S. J.; Lee, D.; Lee, S.; Shin, H. S.; Kim, M. C.; Lee, K. N.; Palmer, D. C.; Theoret, M. R.; Jenkinson, E. J.; Anderson, G.; Restifo, N. P.; Kim, M. G., TSCOT+ thymic epithelial cell-mediated sensitive CD4 tolerance by direct presentation. PLoS Biol 2008, 6 (8), e191.

30. Yang, S. J.; Ahn, S.; Park, C. S.; Choi, S.; Kim, M. G., Identifying subpopulations of thymic epithelial cells by flow cytometry using a new specific thymic epithelial marker, Ly110. J Immunol Methods 2005, 297 (1-2), 265–70.

31. Smedegard, G.; Bjork, J., Sulphasalazine: mechanism of action in rheumatoid arthritis. Br J Rheumatol 1995, 34 Suppl 2, 7–15.

32. Konno, H.; Konno, K.; Barber, G. N., Cyclic dinucleotides trigger ULK1 (ATG1) phosphorylation of STING to prevent sustained innate immune signaling. Cell 2013, 155 (3), 688–98.

33. Zhong, B.; Yang, Y.; Li, S.; Wang, Y. Y.; Li, Y.; Diao, F.; Lei, C.; He, X.; Zhang, L.; Tien, P.; Shu, H. B., The adaptor protein MITA links virus-sensing receptors to IRF3 transcription factor activation. Immunity 2008, 29 (4), 538–50.

34. Zhou, Y.; Fei, M.; Zhang, G.; Liang, W. C.; Lin, W.; Wu, Y.; Piskol, R.; Ridgway, J.; McNamara, E.; Huang, H.; Zhang, J.; Oh, J.; Patel, J. M.; Jakubiak, D.; Lau, J.; Blackwood, B.; Bravo, D. D.; Shi, Y.; Wang, J.; Hu, H. M.; Lee, W. P.; Jesudason, R.; Sangaraju, D.; Modrusan, Z.; Anderson, K. R.; Warming, S.; Roose-Girma, M.; Yan, M., Blockade of the Phagocytic Receptor MerTK on Tumor-Associated Macrophages Enhances P2X7R-Dependent STING Activation by Tumor-Derived cGAMP. Immunity 2020, 52 (2), 357–373 e9.

35. Ohkuri, T.; Kosaka, A.; Ishibashi, K.; Kumai, T.; Hirata, Y.; Ohara, K.; Nagato, T.; Oikawa, K.; Aoki, N.; Harabuchi, Y.; Celis, E.; Kobayashi, H., Intratumoral administration of cGAMP transiently accumulates potent macrophages for anti-tumor immunity at a mouse tumor site. Cancer Immunol Immunother 2017, 66 (6), 705–716.

36. Cheng, N.; Watkins-Schulz, R.; Junkins, R. D.; David, C. N.; Johnson, B. M.; Montgomery, S. A.; Peine, K. J.; Darr, D. B.; Yuan, H.; McKinnon, K. P.; Liu, Q.; Miao, L.; Huang, L.; Bachelder, E. M.; Ainslie, K. M.; Ting, J. P., A nanoparticle-incorporated STING activator enhances antitumor immunity in PD-L1-insensitive models of triple-negative breast cancer. JCI Insight 2018, 3 (22).

37. Jutila, M. A.; Kroese, F. G.; Jutila, K. L.; Stall, A. M.; Fiering, S.; Herzenberg, L. A.; Berg, E. L.; Butcher, E. C., Ly-6C is a monocyte/macrophage and endothelial cell differentiation antigen regulated by interferon-gamma. Eur J Immunol 1988, 18 (11), 1819–26.

38. Marcus, A.; Mao, A. J.; Lensink-Vasan, M.; Wang, L.; Vance, R. E.; Raulet, D. H., Tumor-Derived cGAMP Triggers a STING-Mediated Interferon Response in Non-tumor Cells to Activate the NK Cell Response. Immunity 2018, 49 (4), 754–763 e4.

39. Nicolai, C. J.; Wolf, N.; Chang, I. C.; Kirn, G.; Marcus, A.; Ndubaku, C. O.; McWhirter, S. M.; Raulet, D. H., NK cells mediate clearance of CD8(+) T cell-resistant tumors in response to STING agonists. Sci Immunol 2020, 5 (45).

40. Gilfillan, S.; Ho, E. L.; Cella, M.; Yokoyama, W. M.; Colonna, M., NKG2D recruits two distinct adapters to trigger NK cell activation and costimulation. Nat Immunol 2002, 3 (12), 1150–5.

41. Huntington, N. D.; Vosshenrich, C. A.; Di Santo, J. P., Developmental pathways that generate natural-killer-cell diversity in mice and humans. Nat Rev Immunol 2007, 7 (9), 703–14.

42. Laursen, M. F.; Christensen, E.; Degn, L. L. T.; Jonsson, K.; Jakobsen, M. R.; Agger, R.; Kofod-Olsen, E., CD11c-targeted Delivery of DNA to Dendritic Cells Leads to cGAS- and STING-dependent Maturation. J Immunother 2018, 41 (1), 9–18.

43. Andzinski, L.; Spanier, J.; Kasnitz, N.; Kroger, A.; Jin, L.; Brinkmann, M. M.; Kalinke, U.; Weiss, S.; Jablonska, J.; Lienenklaus, S., Growing tumors induce a local STING dependent Type I IFN response in dendritic cells. Int J Cancer 2016, 139 (6), 1350–7.

44. Genard, G.; Lucas, S.; Michiels, C., Reprogramming of Tumor-Associated Macrophages with Anticancer Therapies: Radiotherapy versus Chemo- and Immunotherapies. Front Immunol 2017, 8, 828.

45. Sinha, P.; Clements, V. K.; Ostrand-Rosenberg, S., Reduction of myeloid-derived suppressor cells and induction of M1 macrophages facilitate the rejection of established metastatic disease. J Immunol 2005, 174 (2), 636–45.

46. Kurahara, H.; Shinchi, H.; Mataki, Y.; Maemura, K.; Noma, H.; Kubo, F.; Sakoda, M.; Ueno, S.; Natsugoe, S.; Takao, S., Significance of M2-polarized tumor-associated macrophage in pancreatic cancer. J Surg Res 2011, 167 (2), e211–9.

47. Porcheray, F.; Viaud, S.; Rimaniol, A. C.; Leone, C.; Samah, B.; Dereuddre-Bosquet,N.; Dormont, D.; Gras, G., Macrophage activation switching: an asset for the resolution of inflammation. Clin Exp Immunol 2005, 142 (3), 481–9.

48. Murray, P. J.; Allen, J. E.; Biswas, S. K.; Fisher, E. A.; Gilroy, D. W.; Goerdt, S.; Gordon, S.; Hamilton, J. A.; Ivashkiv, L. B.; Lawrence, T.; Locati, M.; Mantovani, A.; Martinez, F. O.; Mege, J. L.; Mosser, D. M.; Natoli, G.; Saeij, J. P.; Schultze, J. L.; Shirey, K. A.; Sica, A.; Suttles, J.; Udalova, I.; van Ginderachter, J. A.; Vogel, S. N.; Wynn, T. A., Macrophage activation and polarization: nomenclature and experimental guidelines. Immunity 2014, 41 (1), 14–20.

49. Downey, C. M.; Aghaei, M.; Schwendener, R. A.; Jirik, F. R., DMXAA causes tumor site-specific vascular disruption in murine non-small cell lung cancer, and like the endogenous non-canonical cyclic dinucleotide STING agonist, 2'3'-cGAMP, induces M2 macrophage repolarization. PLoS One 2014, 9 (6), e99988.

50. Zarif, J. C.; Hernandez, J. R.; Verdone, J. E.; Campbell, S. P.; Drake, C. G.; Pienta, K. J., A phased strategy to differentiate human CD14+monocytes into classically and alternatively activated macrophages and dendritic cells. Biotechniques 2016, 61 (1), 33–41.

51. Morehouse, B. R.; Govande, A. A.; Millman, A.; Keszei, A. F. A.; Lowey, B.; Ofir, G.; Shao, S.; Sorek, R.; Kranzusch, P. J., STING cyclic dinucleotide sensing originated in bacteria. Nature 2020, 586 (7829), 429–433.

52. Zhang, X.; Shi, H.; Wu, J.; Zhang, X.; Sun, L.; Chen, C.; Chen, Z. J., Cyclic GMP-AMP containing mixed phosphodiester linkages is an endogenous high-affinity ligand for STING. Mol Cell 2013, 51 (2), 226–35.

53. Diner, E. J.; Burdette, D. L.; Wilson, S. C.; Monroe, K. M.; Kellenberger, C. A.; Hyodo, M.; Hayakawa, Y.; Hammond, M. C.; Vance, R. E., The innate immune DNA sensor cGAS produces a noncanonical cyclic dinucleotide that activates human STING. Cell Rep 2013, 3 (5), 1355–61.

54. Zhou, W.; Whiteley, A. T.; de Oliveira Mann, C. C.; Morehouse, B. R.; Nowak, R. P.; Fischer, E. S.; Gray, N. S.; Mekalanos, J. J.; Kranzusch, P. J., Structure of the Human cGAS-DNA Complex Reveals Enhanced Control of Immune Surveillance. Cell 2018, 174 (2), 300–311 e11.

55. Ahn, J.; Son, S.; Oliveira, S. C.; Barber, G. N., STING-Dependent Signaling Underlies IL-10 Controlled Inflammatory Colitis. Cell Rep 2017, 21 (13), 3873–3884.

56. Davies, B. W.; Bogard, R. W.; Young, T. S.; Mekalanos, J. J., Coordinated regulation of accessory genetic elements produces cyclic di-nucleotides for V. cholerae virulence. Cell 2012, 149 (2), 358–70.

57. Woodward, J. J.; Iavarone, A. T.; Portnoy, D. A., c-di-AMP secreted by intracellular Listeria monocytogenes activates a host type I interferon response. Science 2010, 328 (5986), 1703–5.

58. Corrigan, R. M.; Abbott, J. C.; Burhenne, H.; Kaever, V.; Grundling, A., c-di-AMP is a new second messenger in Staphylococcus aureus with a role in controlling cell size and envelope stress. PLoS Pathog 2011, 7 (9), e1002217.

59. Ryan, R. P.; Fouhy, Y.; Lucey, J. F.; Dow, J. M., Cyclic di-GMP signaling in bacteria: recent advances and new puzzles. J Bacteriol 2006, 188 (24), 8327–34.

60. Gao, D.; Wu, J.; Wu, Y. T.; Du, F.; Aroh, C.; Yan, N.; Sun, L.; Chen, Z. J., Cyclic GMP-AMP synthase is an innate immune sensor of HIV and other retroviruses. Science 2013, 341 (6148), 903–6.

61. Li, X. D.; Wu, J.; Gao, D.; Wang, H.; Sun, L.; Chen, Z. J., Pivotal roles of cGAS-cGAMP signaling in antiviral defense and immune adjuvant effects. Science 2013, 341 (6152), 1390–4.

62. King, K. R.; Aguirre, A. D.; Ye, Y. X.; Sun, Y.; Roh, J. D.; Ng, R. P., Jr.; Kohler, R. H.; Arlauckas, S. P.; Iwamoto, Y.; Savol, A.; Sadreyev, R. I.; Kelly, M.; Fitzgibbons, T. P.; Fitzgerald, K. A.; Mitchison, T.; Libby, P.; Nahrendorf, M.; Weissleder, R., IRF3 and type I interferons fuel a fatal response to myocardial infarction. Nat Med 2017, 23 (12), 1481–1487.

63. Stetson, D. B.; Ko, J. S.; Heidmann, T.; Medzhitov, R., Trex1 prevents cell-intrinsic initiation of autoimmunity. Cell 2008, 134 (4), 587–98.

64. Yang, Y. G.; Lindahl, T.; Barnes, D. E., Trex1 exonuclease degrades ssDNA to prevent chronic checkpoint activation and autoimmune disease. Cell 2007, 131 (5), 873–86.

65. Zhao, Q.; Wei, Y.; Pandol, S. J.; Li, L.; Habtezion, A., STING Signaling Promotes Inflammation in Experimental Acute Pancreatitis. Gastroenterology 2018, 154 (6), 1822–1835 e2.

66. Lan, Y. Y.; Heather, J. M.; Eisenhaure, T.; Garris, C. S.; Lieb, D.; Raychowdhury, R.; Hacohen, N., Extranuclear DNA accumulates in aged cells and contributes to senescence and inflammation. Aging Cell 2019, 18 (2), e12901.

67. Vilalta, M.; Rafat, M.; Giaccia, A. J.; Graves, E. E., Recruitment of circulating breast cancer cells is stimulated by radiotherapy. Cell Rep 2014, 8 (2), 402–9.

68. Gibson, D. G.; Young, L.; Chuang, R. Y.; Venter, J. C.; Hutchison, C. A.3rd,; Smith, H. O., Enzymatic assembly of DNA molecules up to several hundred kilobases. Nat Methods 2009, 6 (5), 343–5.

69. Brinkman, E. K.; Chen, T.; Amendola, M.; van Steensel, B., Easy quantitative assessment of genome editing by sequence trace decomposition. Nucleic Acids Res 2014, 42 (22), e168.

70. Wagner, G. P.; Kin, K.; Lynch, V. J., Measurement of mRNA abundance using RNA-seq data: RPKM measure is inconsistent among samples. Theory Biosci 2012, 131 (4), 281–5.

