## Supplemental Figures for "Human SLC46A2 is the dominant cGAMP importer in extracellular cGAMP-sensing macrophages and monocytes"

Figure S1

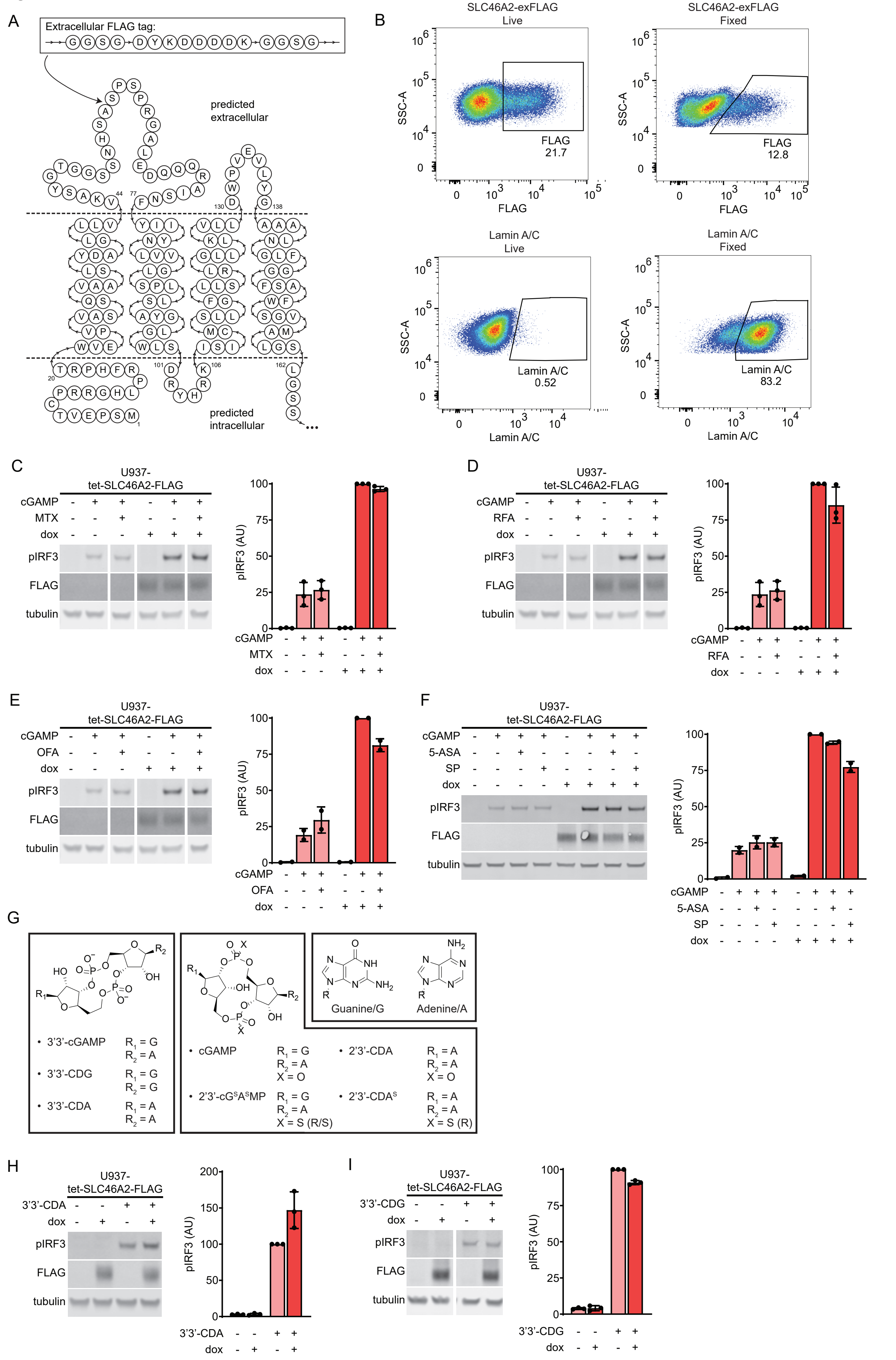
A
Extracellular FLAG tag:
G G S G D Y K D D D D K G G S G
predicted extracellular
predicted intracellular
44
77
130
138
20
101
106
162
1
B
SLC46A2-exFLAG Live
SLC46A2-exFLAG Fixed
SSC-A
FLAG
FLAG 21.7
FLAG 12.8
Lamin A/C Live
Lamin A/C Fixed
Lamin A/C
Lamin A/C 0.52
Lamin A/C 83.2
C
U937-tet-SLC46A2-FLAG
cGAMP
MTX
dox
pIRF3
FLAG
tubulin
pIRF3 (AU)
D
U937-tet-SLC46A2-FLAG
cGAMP
RFA
dox
pIRF3
FLAG
tubulin
pIRF3 (AU)
E
U937-tet-SLC46A2-FLAG
cGAMP
OFA
dox
pIRF3
FLAG
tubulin
pIRF3 (AU)
F
U937-tet-SLC46A2-FLAG
cGAMP
5-ASA
SP
dox
pIRF3
FLAG
tubulin
pIRF3 (AU)
G
3'3'-cGAMP R1 = G R2 = A
3'3'-CDG R1 = G R2 = G
3'3'-CDA R1 = A R2 = A
cGAMP R1 = G R2 = A X = O
2'3'-cGSASMP R1 = G R2 = A X = S (R/S)
2'3'-CDA R1 = A R2 = A X = O
2'3'-CDAS R1 = A R2 = A X = S (R)
Guanine/G
Adenine/A
H
U937-tet-SLC46A2-FLAG
3'3'-CDA
dox
pIRF3
FLAG
tubulin
pIRF3 (AU)
I
U937-tet-SLC46A2-FLAG
3'3'-CDG
dox
pIRF3
FLAG
tubulin
pIRF3 (AU)

Figure S2

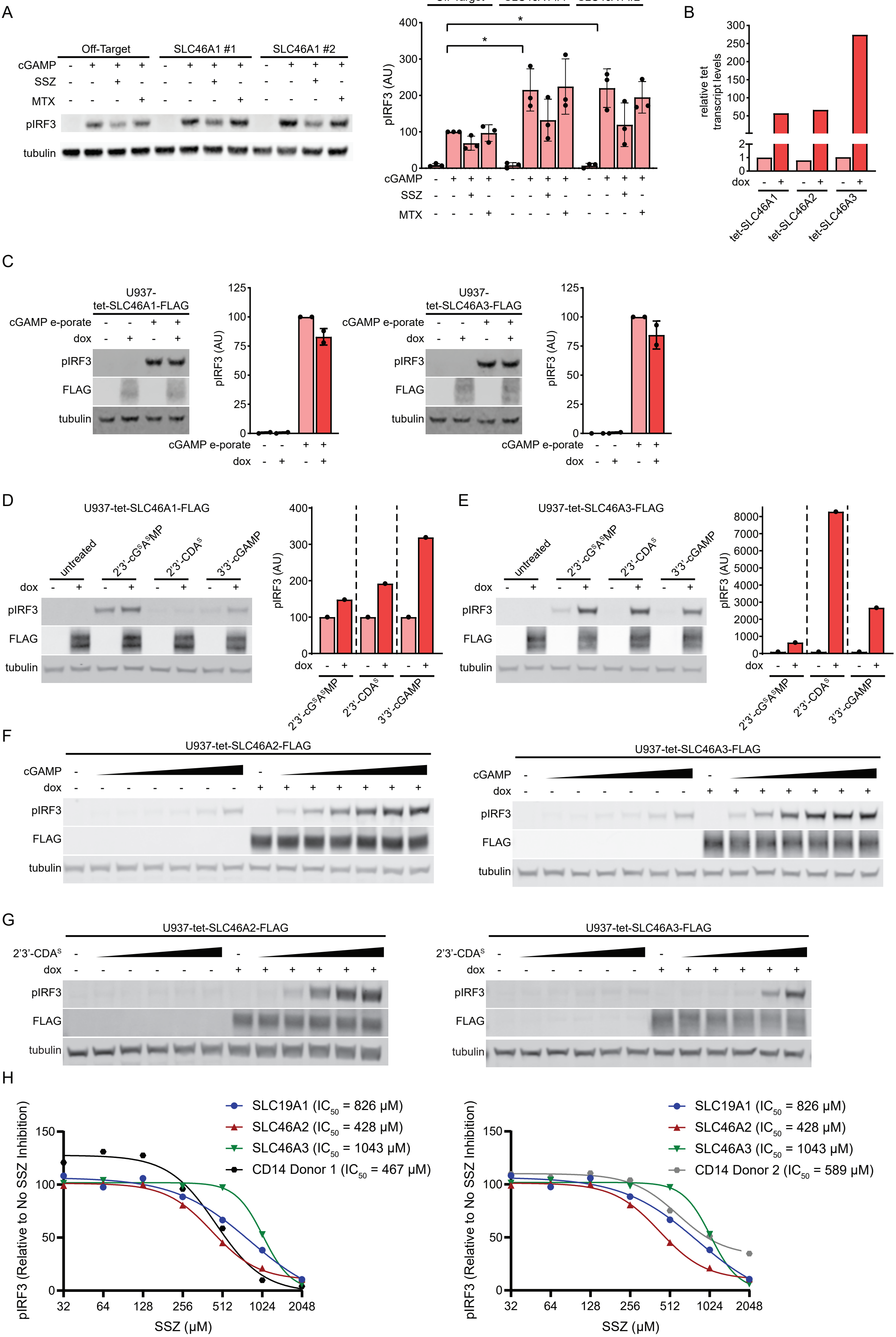
A
Off-Target
SLC46A1 #1
SLC46A1 #2
cGAMP
SSZ
MTX
pIRF3
tubulin
pIRF3 (AU)
B
relative tet transcript levels
dox
tet-SLC46A1
tet-SLC46A2
tet-SLC46A3
C
U937-tet-SLC46A1-FLAG
U937-tet-SLC46A3-FLAG
cGAMP e-porate
dox
FLAG
D
U937-tet-SLC46A1-FLAG
untreated
2'3'-cG^SA^SMP
2'3'-CDA^S
3'3'-cGAMP
E
U937-tet-SLC46A3-FLAG
F
U937-tet-SLC46A2-FLAG
U937-tet-SLC46A3-FLAG
G
2'3'-CDA^S
H
SLC19A1 (IC50 = 826 µM)
SLC46A2 (IC50 = 428 µM)
SLC46A3 (IC50 = 1043 µM)
CD14 Donor 1 (IC50 = 467 µM)
CD14 Donor 2 (IC50 = 589 µM)
pIRF3 (Relative to No SSZ Inhibition)
SSZ (µM)

### Figure S3

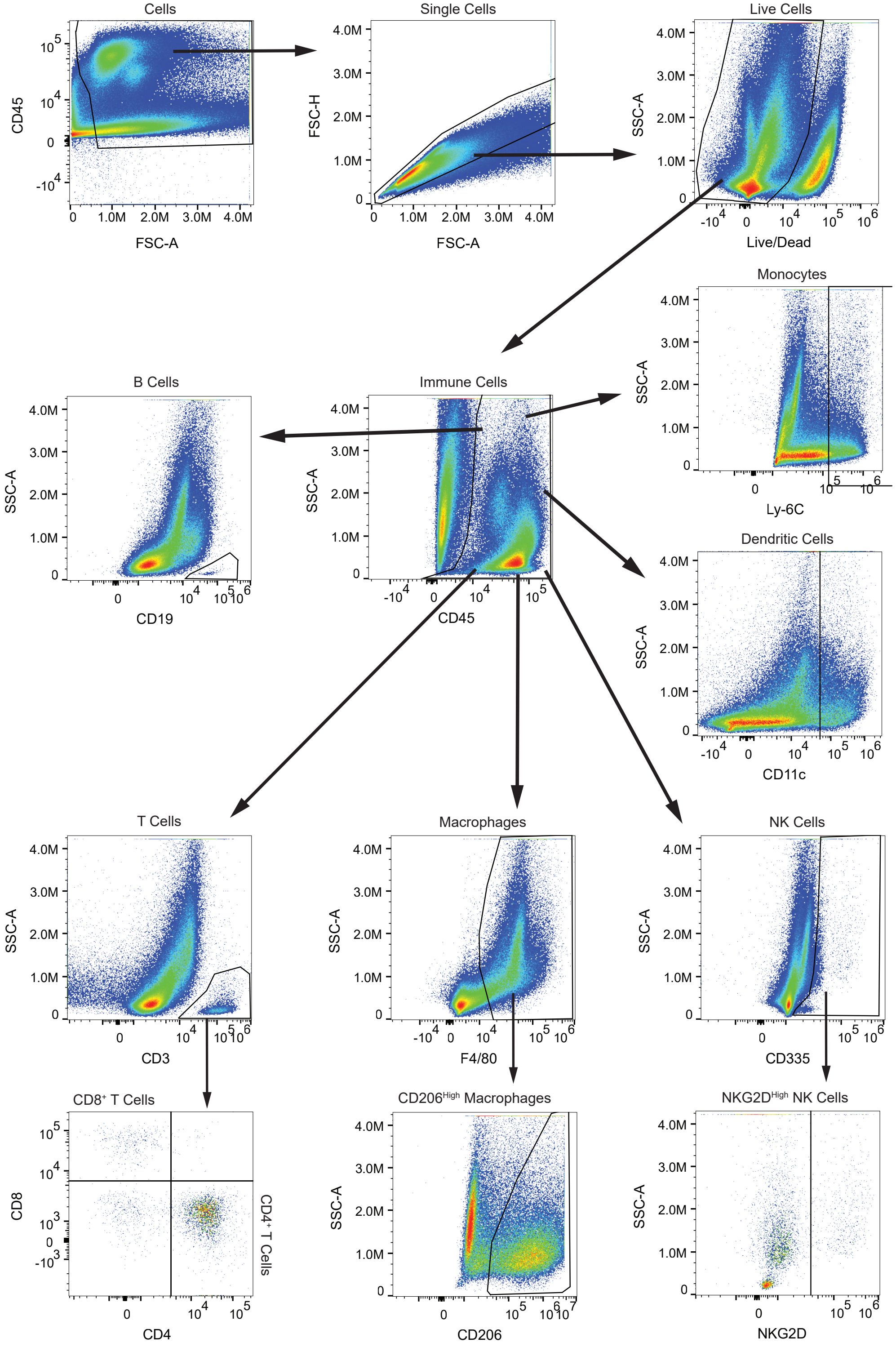
Cells
CD45
FSC-A
Single Cells
FSC-H
FSC-A
Live Cells
SSC-A
Live/Dead
Monocytes
SSC-A
Ly-6C
B Cells
SSC-A
CD19
Immune Cells
SSC-A
CD45
Dendritic Cells
SSC-A
CD11c
T Cells
SSC-A
CD3
Macrophages
SSC-A
F4/80
NK Cells
SSC-A
CD335
CD8⁺ T Cells
CD8
CD4
CD4⁺ T Cells
CD206High Macrophages
SSC-A
CD206
NKG2DHigh NK Cells
SSC-A
NKG2D

### Figure S4

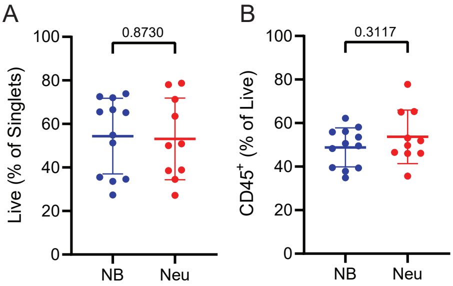


C

| Cell Type | NB STING | Neu STING |
|---|---|---|
| CD45$^+$ (All Immune Cells) | 48.8 | 53.7 |
| CD3$^+$ (All T Cells) | 1.00 | 1.09 |
| CD4$^+$ (Helper T Cells) | 0.74 | 0.82 |
| CD8$^+$ (Cytotoxic T Cells) | 0.09 | 0.10 |
| CD11c$^+$ (Dendritic Cells) | 6.4 | 7.2 |
| CD19$^+$ (B Cells) | 0.15 | 0.14 |
| Ly-6C$^+$ (Monocyte Lineage) | 9.8 | 12.1 |
| F4/80$^+$ (Macrophages) | 18.7 | 22.0 |
| CD335$^+$ (NK Cells) | 0.86 | 0.78 |

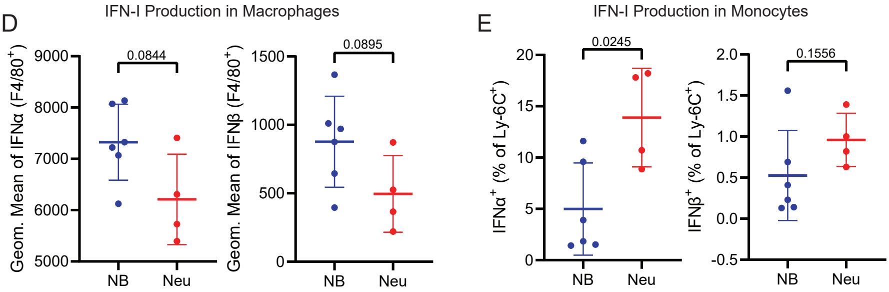

### Figure S5

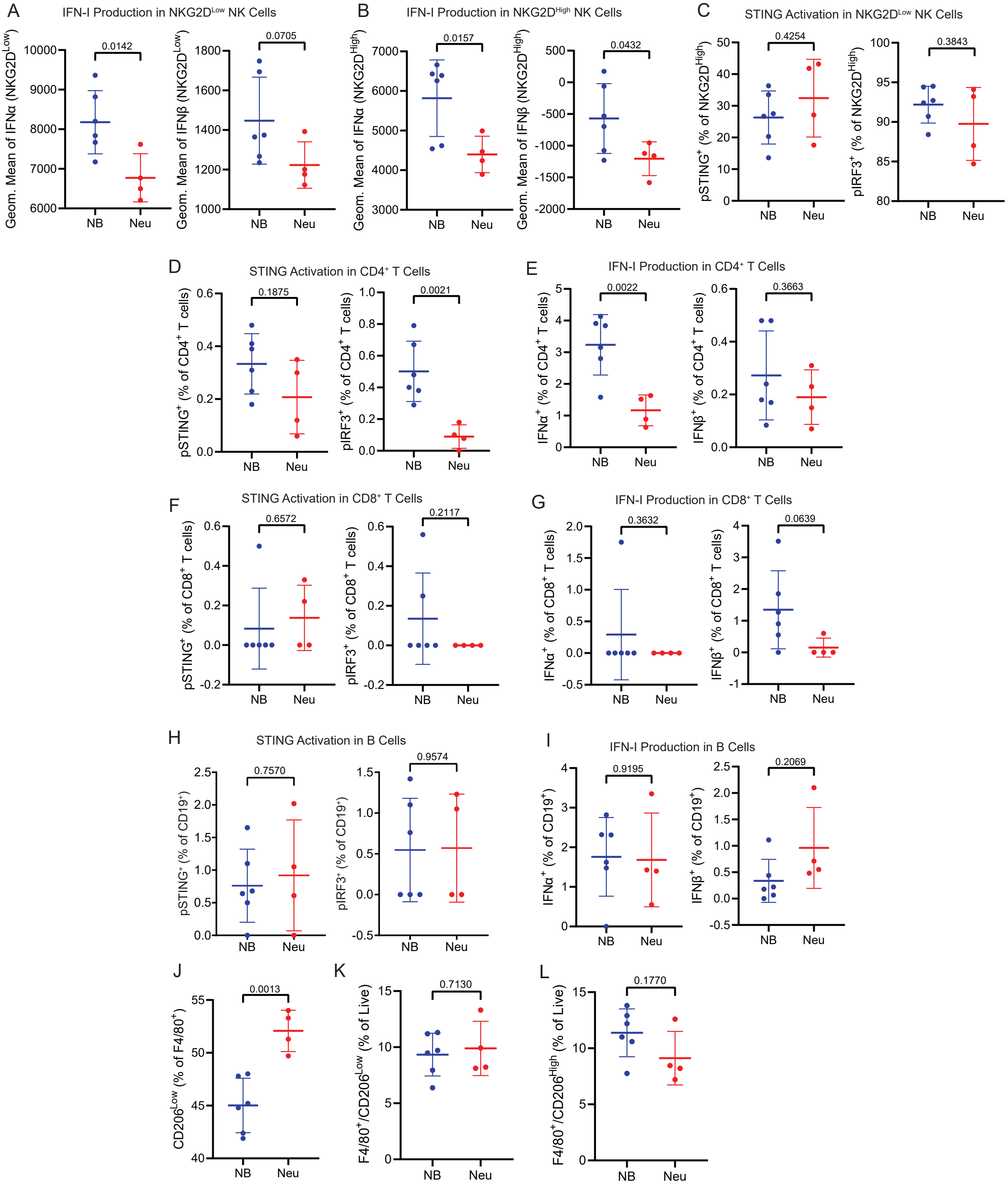
A
IFN-I Production in NKG2D^Low NK Cells
B
IFN-I Production in NKG2D^High NK Cells
C
STING Activation in NKG2D^Low NK Cells
D
STING Activation in CD4⁺ T Cells
E
IFN-I Production in CD4⁺ T Cells
F
STING Activation in CD8⁺ T Cells
G
IFN-I Production in CD8⁺ T Cells
H
STING Activation in B Cells
I
IFN-I Production in B Cells
J
K
L
NB
Neu

Figure S6

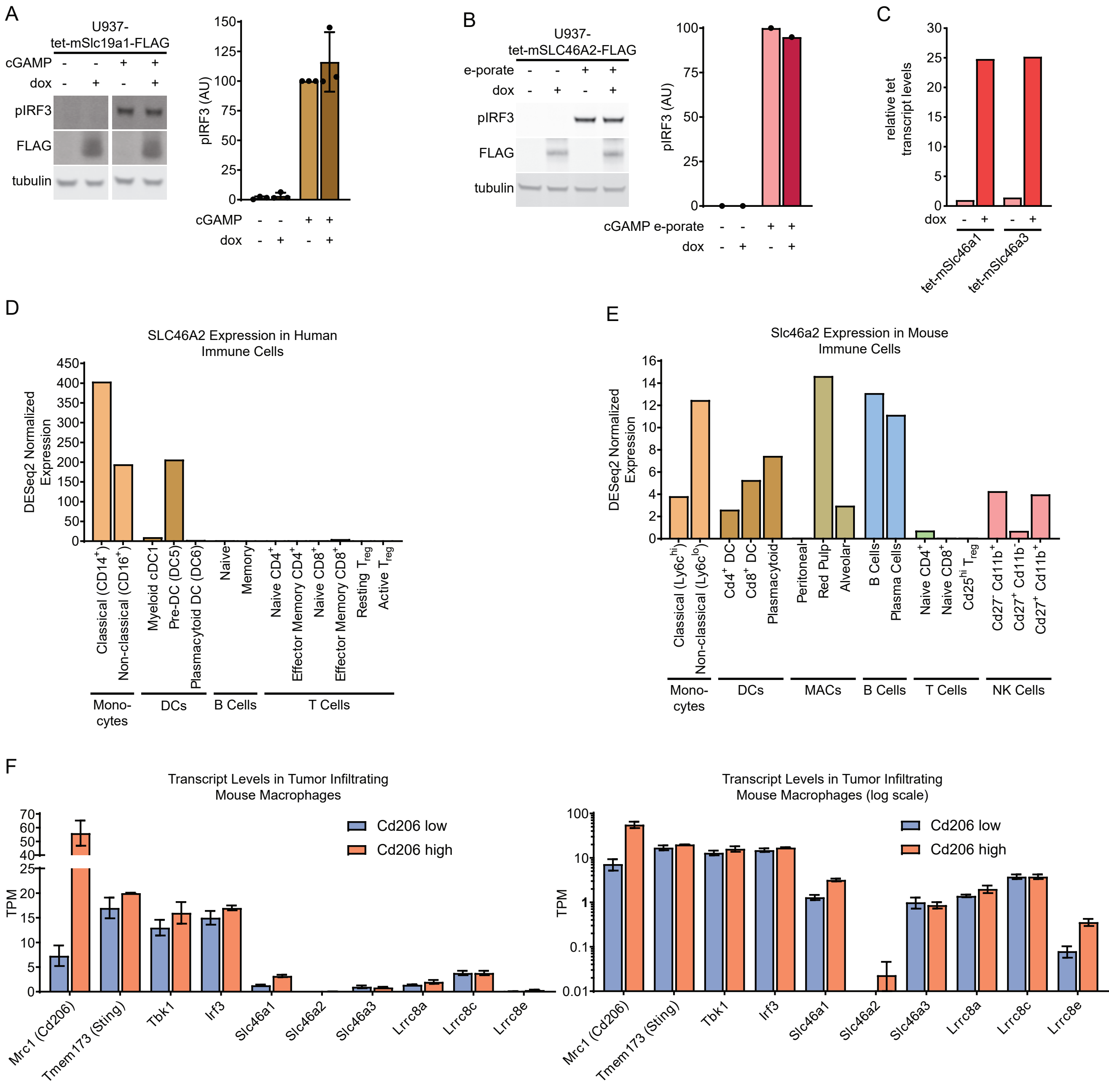
A
U937-
tet-mSlc19a1-FLAG
cGAMP - - + +
dox - + - +
pIRF3
FLAG
tubulin
pIRF3 (AU)
0
25
50
75
100
125
150
cGAMP - - + +
dox - + - +
B
U937-
tet-mSLC46A2-FLAG
e-porate - - + +
dox - + - +
pIRF3
FLAG
tubulin
pIRF3 (AU)
0
25
50
75
100
cGAMP e-porate - - + +
dox - + - +
C
relative tet
transcript levels
0
5
10
15
20
25
30
dox - + - +
tet-mSlc46a1
tet-mSlc46a3
D
SLC46A2 Expression in Human
Immune Cells
DESeq2 Normalized
Expression
0
50
100
150
200
250
300
350
400
450
Classical (CD14$^+$)
Non-classical (CD16$^+$)
Myeloid cDC1
Pre-DC (DC5)
Plasmacytoid DC (DC6)
Naive
Memory
Naive CD4$^+$
Effector Memory CD4$^+$
Naive CD8$^+$
Effector Memory CD8$^+$
Resting $T_{reg}$
Active $T_{reg}$
Mono-
cytes
DCs
B Cells
T Cells
E
Slc46a2 Expression in Mouse
Immune Cells
DESeq2 Normalized
Expression
0
2
4
6
8
10
12
14
16
Classical (Ly6c$^{hi}$)
Non-classical (Ly6c$^{lo}$)
Cd4$^+$ DC
Cd8$^+$ DC
Plasmacytoid
Peritoneal
Red Pulp
Alveolar
B Cells
Plasma Cells
Naive CD4$^+$
Naive CD8$^+$
Cd25$^{hi}$ $T_{reg}$
Cd27$^-$ Cd11b$^+$
Cd27$^+$ Cd11b$^-$
Cd27$^+$ Cd11b$^+$
Mono-
cytes
DCs
MACs
B Cells
T Cells
NK Cells
F
Transcript Levels in Tumor Infiltrating
Mouse Macrophages
TPM
0
5
10
15
20
25
40
50
60
70
Cd206 low
Cd206 high
Mrc1 (Cd206)
Tmem173 (Sting)
Tbk1
Irf3
Slc46a1
Slc46a2
Slc46a3
Lrrc8a
Lrrc8c
Lrrc8e
Transcript Levels in Tumor Infiltrating
Mouse Macrophages (log scale)
TPM
0.01
0.1
1
10
100
Cd206 low
Cd206 high
Mrc1 (Cd206)
Tmem173 (Sting)
Tbk1
Irf3
Slc46a1
Slc46a2
Slc46a3
Lrrc8a
Lrrc8c
Lrrc8e
