## Supplemental Table 1 for "Human SLC46A2 is the dominant cGAMP importer in extracellular cGAMP-sensing macrophages and monocytes"

**Supplemental Table 1. Oligonucleotides used in this study**

| **Oligonucleotides used for cloning:** | |
|---|---|
| **Name** | **Sequence (5'->3')** |
| hSLC19A1_into_dox_fwd | ACCAACTTTCCGTACCACTTCCTACCCTCGTAAAGAATTCAT GGTGCCCTCCAGCC |
| hSLC19A1_into_dox_rev | GTTACTTGTCGTCATCGTCTTTGTAGTCGCCGGATCCGCCC TGGTTCACATTCTGAACACCG |
| mSlc19a1_into_dox_fwd | ACCAACTTTCCGTACCACTTCCTACCCTCGTAAAGAATTCAT GGTGCCCACTGGCCAG |
| mSlc19a1_into_dox_rev | GTTACTTGTCGTCATCGTCTTTGTAGTCGCCGGATCCGCCA GCCTTGGCTTCGACTCTTAAGTC |
| hSLC46A1_into_dox_fwd | ACCAACTTTCCGTACCACTTCCTACCCTCGTAAAGAATTCAT GGAGGGATCTGCCAGCCC |
| hSLC46A1_into_dox_rev | GTTACTTGTCGTCATCGTCTTTGTAGTCGCCGGATCCGCCA GGGGACTGGGGAAACTGCTG |
| mSlc46a1_into_dox_fwd | ACCAACTTTCCGTACCACTTCCTACCCTCGTAAAGAATTCAT GGAAGGAAGAGTTAGTTCAGTAGG |
| mSlc46a1_into_dox_rev | GTTACTTGTCGTCATCGTCTTTGTAGTCGCCGGATCCGCCT GGTGATTGTGGGAATTGTTG |
| hSLC46A2_into_dox_fwd | CGATGTTCCAGATTACGCTGGTGGAGGTGGAGGTTCTAGAA TGAGCCCCGAGGTCACC |
| hSLC46A2_into_dox_rev | TCAGCGGTTTAAACTTAAGCTTGGTACCGAGCTCGGATCCG TGGGTTGCTTAGGTCACCAG |
| mSlc46a2_into_dox_fwd | ACCAACTTTCCGTACCACTTCCTACCCTCGTAAAGAATTCAT GGGTCCAGGGGGCACC |
| mSlc46a2_into_dox_rev | GTTACTTGTCGTCATCGTCTTTGTAGTCGCCGGATCCGCCG CTCCTCTGTTTCTCTGCACACTC |
| hSLC46A3_into_dox_fwd | ACCAACTTTCCGTACCACTTCCTACCCTCGTAAAGAATTCAT GAAGATCCTGTTCGTGGAG |
| hSLC46A3_into_dox_rev | GTTACTTGTCGTCATCGTCTTTGTAGTCGCCGGATCCGCCG CGATCGCTGGCGTCCT |
| mSlc46a3_into_dox_fwd | ACCAACTTTCCGTACCACTTCCTACCCTCGTAAAGAATTCAT GAAAATCAGTTTTATTGAACCAGC |
| mSlc46a3_into_dox_rev | GTTACTTGTCGTCATCGTCTTTGTAGTCGCCGGATCCGCCA TCGCTCGTATGTTCGCTCG |
| GGSG_FLAG_GGSG_top | GGCGGATCCGGCGACTACAAAGACGATGACGACAAGGGCG GATCCGGC |
| GGSG_FLAG_GGSG_bot | GCCGGATCCGCCCTTGTCGTCATCGTCTTTGTAGTCGCCGG ATCCGCC |
| **Primers for qPCR:** | |
| **Name** | **Sequence (5'->3')** |
| tet_3'UTR_fwd | GACAGCCAATGACGGGTAAG |
| tet_3'UTR_rev | CGCCTGTCTTAGGTTGGAGTG |
| actin_fwd | GGCATCCTCACCCTGAAGTA |
| actin_rev | AGAGGCGTACAGGGATAGCA |
| **Primers for genomic sequencing (for measuring CRISPR KO efficiency):** | |
| **Name** | **Sequence (5'->3')** |
| SLC46A2_gene_for | ATCTGCATGTCGCTGCTGG |
| SLC46A2_gene_rev | CACCCAGGAAGCTGGTGATG |

| | |
|---|---|
| SLC46A3_gene_for | AGTCCCATCCACAAGGTATCTC |
| SLC46A3_gene_rev | CATAGCCATTCCTGTCATCGTG |
| **sgRNAs:** | |
| **Name** | **Sequence (5’->3’)** |
| Non-Targeting | GCACTACCAGAGCTAACTCA |
| SLC46A2 | GCCAGACACGGTATCCACGG |
| SLC46A3 | GTTACTATGTCATGTAGTGA |
