## Supplemental Table 2 for "Human SLC46A2 is the dominant cGAMP importer in extracellular cGAMP-sensing macrophages and monocytes"

**Supplemental Table 2 - Antibody Dilutions**

| Antibody | Manufacturer | Dilution |
|---|---|---|
| **For Flow Cytometry** | | |
| Alexa Fluor 594 anti-mouse CD8a Antibody | BioLegend | 1:200 |
| Alexa Fluor® 700 anti-mouse CD45 Antibody | BioLegend | 1:800 |
| Anti-rabbit IgG (H+L), F(ab')2 Fragment (Alexa Fluor 488 Conjugate) | Cell Signaling Technology | 1:500 |
| Anti-rabbit IgG (H+L), F(ab')2 Fragment (Alexa Fluor 647 Conjugate) | Cell Signaling Technology | 1:500 |
| APC anti-mouse F4/80 Antibody | BioLegend | 1:200 |
| Brilliant Violet 510 anti-mouse F4/80 Antibody | BioLegend | 1:400 |
| Brilliant Violet 570 anti-mouse Ly-6C Antibody | BioLegend | 1:200 |
| Brilliant Violet 605 anti-mouse CD335 (NKp46) Antibody | BioLegend | 1:800 |
| Brilliant Violet 650 anti-mouse CD206 (MMR) Antibody | BioLegend | 1:100 |
| Brilliant Violet 785 anti-mouse CD8a Antibody | BioLegend | 1:200 |
| Brilliant Violet 785 anti-mouse CD11c Antibody | BioLegend | 1:400 |
| BUV805 Rat Anti-Mouse CD4, Clone GK1.5 (RUO) | BD Biosciences | 1:200 |
| BV711 Rat Anti-Mouse CD314, Clone CX5 (RUO) | BD Biosciences | 1:400 |
| CD19-PE-Vio700, mouse (DISCONTINUED) | Miltenyi Biotec | 1:100 |
| CD3e Monoclonal Antibody (eBio500A2 (500A2)), PerCP-eFluor 710, eBioscience | Invitrogen | 1:200 |
| DYKDDDDK Tag Antibody (Binds to same epitope as Sigma's Anti-FLAG M2 Antibody) | Cell Signaling Technology | 1:800 |
| FITC anti-mouse CD3ε Antibody | BioLegend | 1:200 |
| FITC Conjugated Anti-Mouse IFN Alpha Antibody, Clone RMMA-1 (MAb) | PBL Assay Science | 1:100 |
| Human TruStain FcX (Fc Receptor Blocking Solution) | BioLegend | 1:100 |
| Lamin A/C (4C11) Mouse mAb (Alexa Fluor 488 Conjugate) | Cell Signaling Technology | 1:100 |
| Mouse Interferon beta (IFN-beta) AssayLite Antibody (APC Conjugate) | AssayPro | 1:25 |
| PE anti-mouse CD11c Antibody | BioLegend | 1:200 |
| PE anti-mouse CD206 (MMR) Antibody | BioLegend | 1:200 |
| PE/Cy7 anti-mouse CD19 Antibody | BioLegend | 1:200 |
| Phospho-IRF-3 (Ser396) (D6O1M) Rabbit mAb (Alexa Fluor 647 Conjugate) | Cell Signaling Technology | 1:50 |
| Phospho-STING (Ser365) (D1C4T) Rabbit mAb | Cell Signaling Technology | 1:200 |
| TruStain FcX (anti-mouse CD16/32) Antibody | BioLegend | 1:100 |
| **For Western Blots** | | |
| α-Tubulin (DM1A) Mouse mAb | Cell Signaling Technology | 1:1000 |
| DYKDDDDK Tag Antibody (Binds to same epitope as Sigma's Anti-FLAG M2 Antibody) | Cell Signaling Technology | 1:1000 |
| IRDye 680RD Goat anti-Mouse IgG (H + L) | LI-COR Biosciences | 1:15000 |

| | | |
|---|---|---|
| IRDye 800CW Goat anti-Rabbit IgG (H + L) | LI-COR Biosciences | 1:15000 |
| Phospho-IRF-3 (Ser396) (D6O1M) Rabbit mAb | Cell Signaling Technology | 1:1000 |
